# From *ATOM* to *GradiATOM*: Cortical gradients support time and space processing as revealed by a meta-analysis of neuroimaging studies

**DOI:** 10.1101/2020.04.29.068411

**Authors:** Giorgia Cona, Martin Wiener, Cristina Scarpazza

**Affiliations:** Department of General Psychology, University of Padua, Via Venezia 8, 35131, Padua, Italy; Padova Neuroscience Center, University of Padua, Italy; Department of Psychology, George Mason University, Fairfax, VA

**Keywords:** space, time, neural, meta-analysis, gradient, ALE method, spatial cognition, temporal processing, timing, fMRI

## Abstract

According to the ATOM (A Theory Of Magnitude), formulated by Walsh more than fifteen years ago, there is a general system of magnitude in the brain that comprises regions, such as the parietal cortex, shared by space, time and other magnitudes (Walsh, 2003).

The present meta-analysis of neuroimaging studies used the Activation Likelihood Estimation (ALE) method in order to determine the set of regions commonly activated in space and time processing and to establish the neural activations specific to each magnitude domain. Following PRISMA guidelines, we included in the analysis a total of 112 and 114 experiments, exploring space and time processing, respectively.

We clearly identified the presence of a system of brain regions commonly recruited in both space and time and that includes: bilateral insula, the pre-supplementary motor area (SMA), the right frontal operculum and the intraparietal sulci. These regions might be the best candidates to form the core magnitude neural system. Surprisingly, along each of these regions but the insula, ALE values progressed in a cortical gradient from time to space. The SMA exhibited an anterior-posterior gradient, with space activating more-anterior regions (i.e., pre-SMA) and time activating more-posterior regions (i.e., SMA-proper). Frontal and parietal regions showed a dorsal-ventral gradient: space is mediated by dorsal frontal and parietal regions, and time recruits ventral frontal and parietal regions.

Our study supports but also expands the ATOM theory. Therefore, we here re-named it the ‘*GradiATOM*’ theory (Gradient Theory of Magnitude), proposing that gradient organization can facilitate the transformations and integrations of magnitude representations by allowing space- and time-related neural populations to interact with each other over minimal distances.

## Introduction

> *“But Einstein came along and took space and time out of the realm of stationary things and put them in the realm of relativity… because time and space are modes by which we think and not conditions in which we live.”*
>
> — Dimitri Marianoff, Einstein: An intimate study of a great man

Imagine the following scenario. You went on a safari in the savanna. Unexpectedly, you find yourself close to an angry and hungry lion: the lion wishes to catch you while you try to evade capture. If you want to survive you need to choose the best strategy to reach the safest place in the shortest possible time, so you need to estimate space and time jointly and accurately. This example, although extreme, emphasizes how it is (and has been) essential for humans to form a representation that jointly contains both spatial and temporal information in order to survive and, in broader terms, to face evolutionary challenges (Bufacchi & Iannetti, 2018). Time and space have indeed a close relationship in human perception, representation and action as providing a natural framework for organizing our behaviour and experience.

Behavioural data have hinted at such interrelations between the processing of these magnitudes, showing a reciprocal influence – and interference – between space and time perception in both monkeys and humans (Casasanto & Boroditsky, 2008; Merritt, Casasanto, & Brannon, 2010; Xuan, Zhang, He, & Chen, 2007). For example, the judgement of stimulus length is affected by its concurrent duration, and *vice versa* (Cai & Connell, 2015), or saccadic eye movements compress the judgments of both spatial and temporal magnitude (Morrone, Ross, & Burr, 2005), and prism adaptations aimed to produce a spatial manipulation can cause misperceptions of temporal intervals (Magnani, Oliveri, Mancuso, Galante, & Frassinetti, 2011; Oliveri, Magnani, Filipelli, Avanzi, & Frassinetti, 2013). Further, a well-known interaction between location and time is the Spatial Temporal Association of Response Codes, or STEARC effect (Bonato, Zorzi, & Umilta, 2012; Ishihara, Keller, Rossetti, & Prinz, 2008), which reflects the tendency to associate horizontal locations with the concepts of past versus future, or before versus after (Ishihara et al., 2008; Santiago, Neto Dde, Gandini, & Tabak, 2008; Torralbo, Santiago, & Lupianez, 2006; Vallesi, Weisblatt, Semenza, & Shaki, 2014; Weger & Pratt, 2008).

In addition to behavioural findings, several lines of neuropsychological and neuroimaging research point out the idea that similar neurocognitive systems support processing in the two domains. For example, damage to the right parietal cortex often leads to hemispatial neglect, which results in deficits in directing visuospatial attention to the part of the space contralateral to the lesion. Critically, left neglect patients often show selective “neglect” of temporal representation, exhibiting – for example – slower performance for item occurring before a time-related reference (Bonato, Saj, & Vuilleumier, 2016). Likewise, disruptions following transcranial magnetic stimulation – TMS (Oliveri et al., 2009; Riemer, Diersch, Bublatzky, & Wolbers, 2016), and neuroimaging investigated with fMRI (Peer, Salomon, Goldberg, Blanke, & Arzy, 2015), PET (Coull & Nobre, 1998), structural imaging (Thiebaut de Schotten et al., 2011) and EEG studies (Vallesi, McIntosh, & Stuss, 2011) consistently revealed partial overlaps between the brain systems involved in time and space processing, in particular in the parietal cortex. A variety of fMRI studies have consistently shown that time and space, together with other magnitudes, such as number, size etc., share common activations in the parietal cortex, which has been suggested as the best possible candidate for the locus of magnitudes processing (Dormal, Dormal, Joassin, & Pesenti, 2012; Hayashi et al., 2013; Skagerlund, Karlsson, & Traff, 2016). This line of research is well represented and integrated in “A Theory Of Magnitude” –ATOM– (Bueti & Walsh, 2009; Walsh, 2003). According to ATOM, space, time, and numbers would share a common system of processing and neuroanatomical structures, likely located in neurons of the parietal cortex for evolutionary reasons, as they would be in the service of prompt sensorimotor transformations and actions. Importantly, beyond the parietal cortex, possible overlaps in other brain regions, especially over frontal regions, are far from being clearly established (Coull, Charras, Donadieu, Droit-Volet, & Vidal, 2015; Li, Chen, Han, Chui, & Wu, 2012).

Although several findings support such abstract, common, neural representation of magnitudes, other patterns of findings seem to contradict this view. Some patients, indeed, have reported to show disorientation limited to one single magnitude domain (Aguirre & D’Esposito, 1999). Alzheimer’s patients, for example, typically lose first orientation in time, and only later in space (Daniel, Crovitz, & Weiner, 1987). Also, behavioural studies revealed an asymmetry in the interference between spatial and temporal perceptions suggesting mechanisms that, even if interacting with each other, are separate rather than a common representation of magnitude (Casasanto, Fotakopoulou, & Boroditsky, 2010; Merritt et al., 2010). Interference effects between dimensions, as predicted by ATOM, appear to be highly context dependent and inconstant. For example, while larger stimuli engender longer perceptions of time, the converse is not observed, with longer times having no influence on the perception of size (Riemer, Trojan, Kleinbohl, & Holzl, 2012).

### Neural activations in the space and time domains

*Space-related activations.* In a previous meta-analysis (Cona & Scarpazza, 2019), we analysed neuroimaging data obtained from those studies that explored spatial processing in a variety of cognitive functions. We found a consistent activation in fronto-parietal regions belonging to the Dorsal Attention Network (DAN). Based on previous literature, representational and attentional mechanisms are assigned to the DAN, with fronto-parietal regions being implied in spatial representational processes as well as in allocating attentional resources to both external visuospatial stimuli and internal representations (Luckmann, Jacobs, & Sack, 2014). A large body of evidence suggests indeed the presence of topographic maps in the dorsal frontoparietal regions (Jerde & Curtis, 2013; Serences & Yantis, 2007; Szczepanski, Konen, & Kastner, 2010). Maps of space were also discovered in a region close to the frontal operculum (Hagler & Sereno, 2006).

The frontal operculum and the anterior insular cortex of both hemispheres were shown to be consistently activated in the meta-analysis of spatial tasks (Cona & Scarpazza, 2019) and are supposed to be involved in dynamically prioritizing the topographical maps formed in frontal and parietal regions on the basis of the external stimuli’s saliency and the internal, top-down rules and goals (Serences & Yantis, 2007). Consistent activation of SMA, and mostly pre-SMA, across spatial studies was also found and was shown to reflect hierarchical accumulation and sequential integration of information into higher-order spatial representations (Bahlmann et al., 2009; Cona & Semenza, 2017).

#### Time-related activations

Several meta-analyses were run in order to provide a comprehensive overview of the neural substrates of temporal processing. Wiener et al., 2010 performed a meta-analysis of neuroimaging data to explore the influence of the type of task response (perceptual vs. motor) and the duration of the temporal stimuli (sub-vs. supra-second) in the neural bases of temporal processing. Sub-second timing durations were shown to mainly activate sub-cortical networks, including the basal ganglia and cerebellum, whereas supra-second durations were more associated with cortical regions, such as the prefrontal cortex and SMA. Interestingly, a functional gradient was found in the SMA regions, wherein motor tasks are more likely to activate SMA proper, whereas perceptual tasks are associated with the pre-SMA activations (see also Wiener et al., 2011 and Schwartze et al., 2012). Importantly, this meta-analysis demonstrated that the only structures consistently activated across all timing conditions were the SMA and the right inferior frontal gyrus (Wiener et al., 2010).

Two very recent meta-analyses (Nani et al., 2019; Teghil et al., 2019) further explored the differential involvement of brain structures in relation to the nature of the timing task conditions.

The study by Teghil et al., 2019 (Teghil et al., 2019), in particular, showed that the core timing network, shared across all the timing task conditions, involves SMA inferior frontal gyrus and insula, intraparietal sulcus and basal ganglia, and that these regions were preferentially more active when attending to external stimuli for timing, rather than internally cued timing. Separately, the meta-analysis by Nani et al., 2019 additionally supported previous findings for separable networks across different timing contexts, but with the SMA as the only “core” region active across all of them.

### The present study: The Gradient Hypothesis

Here, we present a quantitative, activation likelihood estimation (ALE) meta-analysis of neuroimaging data on space and time domains, designed to determine both the common and the specific activations underlying the two domains, and further provide insight into their organizational structure. The ALE method treats activated foci of brain regions as three-dimensional Gaussian probability distributions centered at the given coordinates (Eickhoff et al., 2009; Laird et al., 2005). Two maps of ALE distributions are obtained, for space and time domains respectively, and are statistically thresholded. Three possible scenarios regarding the organizational overlap between time and space maps can be hypothesized, as represented in Figure 1.

**Figure 1.**
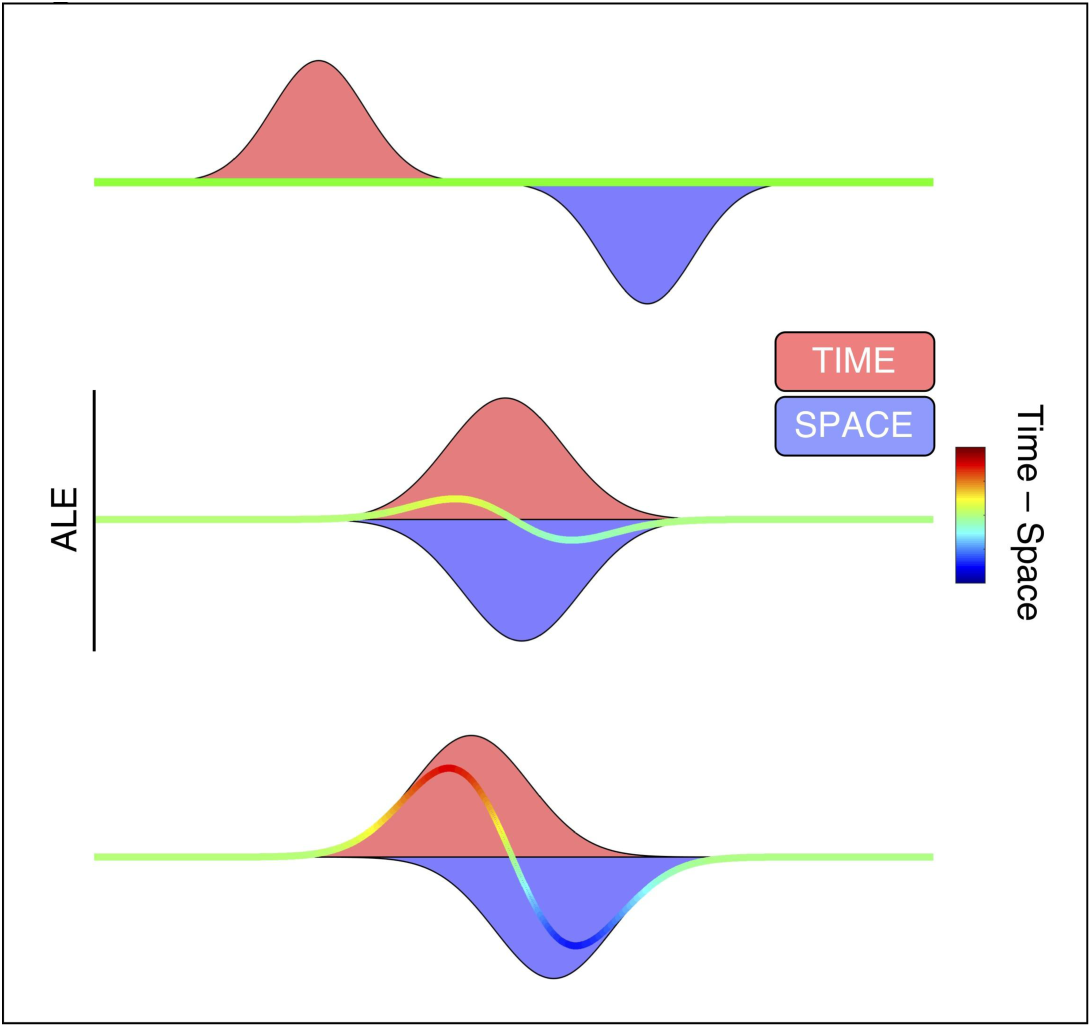
Three hypotheses related to the potential overlap of time and space representations in the brain. In all three hypotheses, Time and Space ALE scores are expressed as positive and negative values, respectively. Top: If Time and Space involve separate brain regions, then combining both representations will not result in any overlap between the two. Middle: If Time and Space involve the same regions, with only slight variation between them, then combining both will lead to an area of full overlap with only marginal differences between the two maps. Bottom: If Time and Space involve partially overlapping brain regions, then the combination of both will result in a transition zone from one dimension to the other; this transition should be expressed as a gradient of activation likelihood.

If activations related to space and time processing are fully separated, we should expect that the related ALE maps of distributions will not overlap at all and the difference in the subtraction analysis, which consists of subtracting the activation of one (e.g., space) to the other domain-related distribution (e.g., time), is flat (in the Figure 1, the green line in the middle). On the other hand, if space and time domains share a complete overlap in their activations, we should observe that time and space ALE maps are in turn nearly fully overlapping. This would lead to absence of domain-related activated clusters of voxels in the subtraction analysis (and so the difference would show only a slight bump in either direction). The third possible scenario is that space and time domains consist of partially overlapping representations, resulting in a partial overlap of the two ALE maps. In the subtraction analysis, this would be reflected in a distribution of domain-related activated clusters along a gradient from one dimension to the other (the red-to-blue transition in Figure 1). In this scenario, a transition between spatial and temporal processing would thus occur in a “gradiential” manner along adjacent cortical regions.

A gradient is an axis of variance in structural and/or functional cortical features, along which brain areas are situated in a spatially contiguous order; areas that resemble each other in relation to those features fall in closer positions along the gradient. The idea of gradients in the brain is relatively new but is gaining increasing attention among researchers (Huntenberg, Bazin & Margulies, 2018). The spatial organization of cortical areas is indeed not arbitrary. Recent evidence shows that gradients are a fundamental, universal, organizing principle whereby brain areas are situated along a hierarchy of continuous gradients for sensory, motor and cognitive interactions (Huntenburg et al., 2018; Margulies et al., 2016). Hierarchical gradients emerge from unimodal sensory areas and motor areas and radiate toward higher-order areas in the temporal, parietal and prefrontal cortex, culminating in the default mode network and salience network (Margulies et al., 2016; Vázquez-Rodríguez et al., 2019). Spatial gradients serve increasingly and hierarchically abstract orders of representation along an axis that separate concrete (perceptual and physical) categories in sensorimotor regions from more abstract concepts encoded in transmodal areas (Huth et al., 2012). Along this hierarchy, low-level sensory features are increasingly abstracted and integrated with concepts from other systems.

A gradient transition of time and space representation appears the most likely scenario on the basis of the ATOM view, which points to a close proximity in the brain so that space and time metrics can be easily coupled in the service of human perception, representation and action (Bueti & Walsh, 2009). As such, the present study represents the first attempt to explore the ‘gradient hypothesis’ with respect to the space and time domains.

## Materials and methods

### Studies selection

For the space domain, we selected the studies already included in our previous meta-analysis following the inclusion criteria described below (Cona & Scarpazza, 2019), whereas for the time processing we conducted a new in-depth search up to February 2019. More specifically, regarding space processing, two hundred and ninety possible eligible papers were identified through database search and additional 167 studies were found by means of the “related articles” function of the PubMed database and by tracing the references from review articles and the identified papers. This yielded to an initial identification of 457 papers. Regarding time processing, five hundred and ninety-five possible eligible papers were identified through database search and additional twenty-seven studies were found by means of the “related articles” function of the PubMed database and by tracing the references from review articles and the identified papers. This yielded an initial identification of 622 papers.

Studies that met the following inclusion criteria have been included in the current research:

i. studies using functional magnetic resonance imaging (fMRI) or positron emission tomography (PET);
ii. studies analyzing the data using univariate approach that revealed localized increased activation (i.e. studies using machine learning and multivoxel pattern analysis were excluded; studies analyzing the data using functional connectivity or related techniques have been discharged);
iii. studies performed a whole brain analysis (i.e. articles that performed only region of interest (ROI) or small volume correction (SVM) analysis have been excluded);
iv. studies that are peer-reviewed articles reporting novel data on the spatial processing in healthy individuals;
v. studies that report a clear higher activation during spatial or temporal processing compared with a control condition;
vi. studies that used a task clearly linked to spatial processing or temporal processing (e.g. studies using mixed task, for example involving both space and time, as for example (Formisano et al., 2002), were excluded);
vii. studies that did not focus on isolating specific brain regions’ activations (e.g., for example, studies on navigation and long-term memory aimed at detecting parahippocampal gyrus and posterior cingulate cortex activation);
viii. studies including more than 5 participants;
ix. studies that report results in a standardized coordinate space (e.g. (Talairach & Tournoux, 1988), or Montreal Neurologic Institute –MNI).

### Systematic Review

The literature screening and final selection has been performed according to the PRISMA guidelines (Liberati et al., 2009; Moher, Liberati, Tetzlaff, Altman, & Group, 2009). This procedure is summarized in the PRISMA flow diagrams that are available within the files: “Supplementary Information A and B” (for space and time, respectively). Applying the PRISMA procedure, a total of 110 original articles were found eligible to be included in the systematic review on space processing and 110 original articles were found eligible to be included in the systematic review on time processing (see file “Supplementary Information C and D” for the list of the studies included). One author (CS) and a student (NT, in the acknowledgements) extracted and checked the data independently. Two additional authors (CG and MW) double-checked random data and also double-checked data in case of discordance between the first two extractions. Two databases (one for space and one for time) were created with the following features of each study: the number of subjects, the specific task used, the contrast performed, the coordinate system, the coordinate localization (brain regions), the p value criteria (corrected, uncorrected) and the associated statistic (t value, z score). In order to avoid dependency across experiment maps that might negatively impact on the validity of the meta-analysis results, for each included study only the contrast that most strongly reflected the process that the current meta-analysis aimed to investigate has been selected, in line with the recent meta-analysis guidelines (Muller et al., 2018). Two out of 110 studies included in the meta-analysis on space processing (Jordan, Wustenberg, Heinze, Peters, & Jancke, 2002; Seurinck, Vingerhoets, de Lange, & Achten, 2004) performed the same experiment using two independent samples as they analyzed males and females separately. As a consequence, two independent contrasts were selected from these studies without the need to adjust for multiple contrasts. This procedure led to the inclusion in the meta-analysis of 110 studies, resulting in 112 experiments, with “study” referring to a paper, and “experiment” referring to an individual contrast reported in each paper. Similarly, four out of the 110 studies included in the meta-analysis on time processing (Chen, Penhune, & Zatorre, 2008; Coull & Nobre, 1998; Gandour et al., 2002; Hayashi et al., 2015) performed the same experiment using two independent samples. As a consequence, four independent contrasts were selected from these papers without the need to adjust for multiple contrasts. This procedure led to the inclusion in the meta-analysis of 110 studies, resulting in 114 experiments.

### The meta-analysis

The current study followed the most recent guidelines for the meta-analysis (Muller et al., 2018). Talairach coordinates were reported into MNI space before performing the meta-analysis using a linear transformation (Laird et al., 2010; Lancaster et al., 2007). For a quantitative assessment of inter study convergence the Activation Likelihood Estimation (ALE) method (Eickhoff et al., 2009; Laird et al., 2005; Turkeltaub, Eden, Jones, & Zeffiro, 2002) has been applied. The peaks of enhanced activation during spatial (or temporal) processing compared to the control condition were used to generate an ALE map, using the revised ALE algorithm (Turkeltaub et al., 2012) running under Ginger ALE software (http://brainmap.org/ale/) version 3.0.2. This approach aims to identify areas with a convergence of reported coordinates across experiments that is higher than expected from a random distribution of foci. Briefly, this algorithm treats activated foci of brain regions as three-dimensional Gaussian probability distributions centered at the given coordinates (Eickhoff et al., 2009; Laird et al., 2005). The algorithm incorporates the size of the probability distributions by considering the sample size of each study. Moreover, the algorithm utilizes the random-effect rather than the fixed-effect inference. It does so by testing the above chance clustering between contrasts rather than the above-chance clustering between foci. Inference is then sought regarding regions where the likelihood of activation being reported in a particular set of experiments is higher than expected by chance, i.e., where there is a non-random convergence. For further details on the ALE method please refer to the original publications (Eickhoff, Bzdok, Laird, Kurth, & Fox, 2012; Eickhoff et al., 2009; Turkeltaub et al., 2012). The checklist for neuroimaging meta-analysis is available within the file “Supplementary Information E”.

To investigate the neural activations respectively associated with space and time, two separate meta-analyses were run. Statistical ALE maps were thresholded using cluster level FWE correction at p<0.05 (cluster-forming threshold at voxel-level P<0.001) (Eickhoff et al., 2016) in line with the recent guidelines for coordinate based meta-analysis (Muller et al., 2018).

Furthermore, to explore brain regions that are specifically activated in space as compared with time, and vice versa, a discriminability (i.e., subtraction) analysis was run between the ALE maps of space and time. This procedure allows one to test if two sets of foci (i.e., space and time) statistically differ in spatial convergence. To perform the discriminability analysis, the experiments contributing to either analysis (space and time) were pooled together and then, recursively for 10,000 times, randomly divided into two groups of the same size as the original sets of data (Eickhoff et al., 2011). An empirical null distribution of ALE-score differences between the two conditions was created subtracting, for each of the 10.000 permutation, the voxelwise ALE scores of these two randomly assembled sets of foci from one another. The true results were then compared with the null distribution. Based on this permutation procedure, the map of true differences was then thresholded using a corrected p<0.05 and an extent threshold of 100 voxels was applied to eliminate minor, presumably incidental, findings. To simplify interpretation of ALE contrast images, they are converted to Z scores to show their significance instead of a direct ALE subtraction. This discriminability analysis yielded three different outputs: brain regions that are specifically activated for space as compared to time (space ALE maps > time ALE maps); brain regions that are specifically activated for time as compared to space (time ALE maps > space ALE maps); brain regions that are similarly activated by the two domains (conjunction analysis between space and time ALE maps).

## Results

### Activations related to space processing

*Space-related activations.* The meta-analysis of all the studies exploring spatial processing included 1333 foci from 112 experiments for a total of 1544 participants (see Table 1, Figure 2a). The minimum cluster size for the cluster to be considered statistically significant was 1232 mm3. The results showed high areas of convergence in a large cluster (41680 voxels) extended within the bilateral dorsal parietal regions (BA 7) including precunei, superior parietal lobules and the regions surrounding the intraparietal sulci, and superior and middle occipital cortices. Furthermore, two large clusters of significant convergence were found in the middle and inferior frontal cortices and in dorsal frontal regions including bilateral frontal eye field (FEF) (16144 and 14880 voxels in the right and left hemisphere, respectively). Additional clusters were located in the pre-SMA (9520 voxels) and in the insulae, bilaterally (2832 and 3768 voxels in the right and left hemispheres respectively).

**Table 1.** Significant activation likelihood clusters for the analysis of space processing.

| Cluster |  |  |  |
| --- | --- | --- | --- |
| Cluster | Coordinates | Brain Region | Broadman Area |
| 49832 | 18 -64 56 | Precuneus | 7 |
|  | -38 -44 44 | Superior Parietal Lobule | 7 |
|  | -16 -66 54 | Precuneus | 7 |
|  | 32 -48 52 | Superior Parietal Lobule | 7 |
|  | 40 -42 44 | Inferior Parietal Lobule | 40 |
|  | -24 -56 58 | Superior Parietal Lobule | 7 |
|  | 28 -66 42 | Superior Occipital Gyrus | 39 |
|  | 34 -84 16 | Middle Occipital Gyrus | 19 |
|  | 36 -78 32 | Superior Occipital Gyrus | 39 |
|  | -28 -82 24 | Middle Occipital Gyrus | 19 |
|  | 26 -52 66 | Superior Parietal Lobule | 7 |
|  | 48 -60 -4 | Fusiform Gyrus | 37 |
|  | 24 -60 24 | Precuneus | 31 |
|  | -30 -86 34 | Middle Occipital Gyrus | 19 |
|  | 44 -82 4 | Middle Occipital Gyrus | 19 |
|  | 50 -68 6 | Middle Occipital Gyrus | 19 |
|  | -22 -70 30 | Superior Occipital Gyrus | 7 |
|  | 56 -60 -12 | Fusiform Gyrus | 37 |
| 16144 | 28 -4 56 | Middle Frontal Gyrus | 6 |
|  | 52 10 24 | Inferior Frontal Gyrus | 44 |
|  | 46 6 42 | Precentral Gyrus/FEF | 8 |
| 14880 | -28 2 54 | Middle Frontal Gyrus | 6 |
|  | -50 12 28 | Inferior Frontal Gyrus | 44 |
|  | -40 -14 56 | Precentral Gyrus/FEF | 8 |
|  | -40 30 26 | Middle Frontal Gyrus | 9 |
| 9520 | 2 14 50 | Pre-SMA | 6 |
|  | 8 28 36 | Pre-SMA | 8 |
| 3768 | -32 24 -4 | Insula | 13 |
| 3592 | -46 -68 -8 | Inferior Occipital Gyrus | 19 |
|  | -48 -72 8 | Middle Occipital Gyrus | 19 |
| 2832 | 34 22 2 | Insula | 13 |

**Figure 2.**
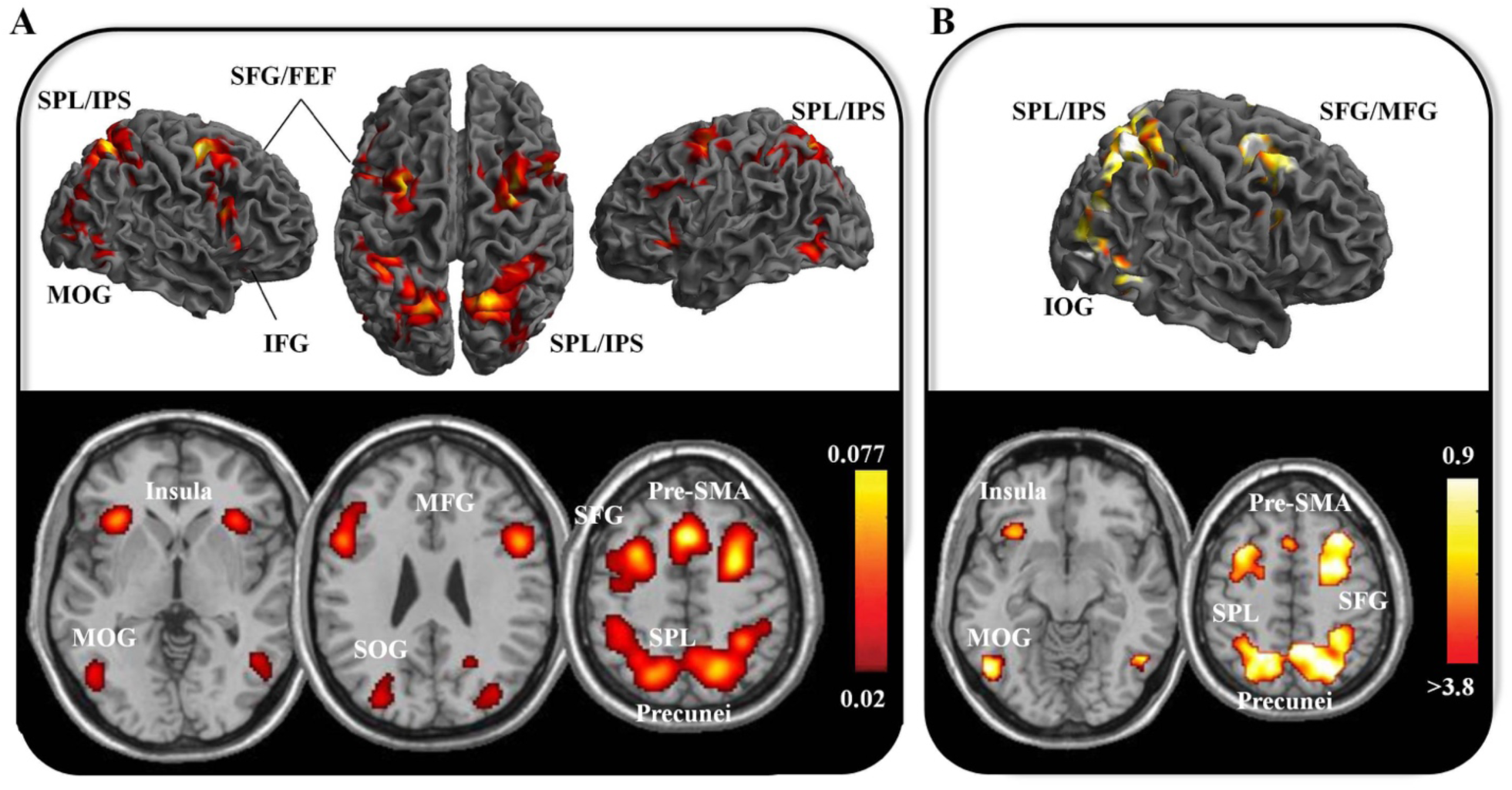
Space related brain activations. A) Brain regions that are activated during tasks requiring space processing. Colors indicate the ALE values for each voxel above the threshold (where yellow indicates the most significant ALE values); B) Brain regions where convergence in the coordinates of activation among the studies is higher during tasks requiring processing of space rather than time (Space > Time). To simplify the interpretation of ALE contrast images, they are converted to z scores to show their significance instead of a direct ALE subtraction (z-score for each voxel above the threshold). SPL = Superior Parietal Lobule; IPS = IntraParietal Sulcus; IFG = Inferior Frontal Gyrus; MFG = Middle Frontal Gyrus; SFG = Superior Frontal Gyrus; FEF = Frontal Eye Field; pre-SMA = pre-Supplementary Motor Area; IOG = Inferior Occipital Gyrus; MOG = Middle Occipital Gyrus; SFG = Superior Occipital Gyrus.

### Space minus time related activations

A meta-analysis that identified brain activations that are more consistently activated for space compared to time (Space > Time direct contrast) was then run. This meta-analysis provided the identification of activations that were specific for space processing (see Table 2, Figure 2b). Strong areas of convergence were still found in the bilateral dorsal fronto-parietal regions and in occipital regions. The biggest cluster of consistent activations (41680 voxels) was detected bilaterally in the precuneus extending to the superior parietal lobules and to the superior occipital cortices. Additional clusters were found in the dorsal frontal regions including superior and middle frontal cortex, bilaterally (9568 voxels and 6152 voxels in the right and left hemisphere, respectively), and in the left inferior frontal gyrus (3128 voxels). Smaller clusters were also found in the pre-SMA (976 voxels) and in the left insula (736 voxels).

**Table 2.** Significant activation likelihood clusters for the space > time analysis.

| **Cluster** | **Coordinates** | **Brain Region** | **Broadman Area** |
| --- | --- | --- | --- |
| 41680 | 6 -65 48 | Precuneus | 7 |
|  | -4 -65 47 | Precuneus | 7 |
|  | -27 -45 50 | Superior Parietal Lobule | 7 |
|  | -35 -42 44 | Superior Parietal Lobule | 7 |
|  | 26 -86 20 | Middle Occipital Gyrus | 19 |
|  | 40 -86 16 | Middle Occipital Gyrus | 19 |
|  | -54 -30 40 | Superior Parietal Lobule | 7 |
|  | -24 -68 26 | Superior Occipital Gyrus | 39 |
|  | 32 -44 42 | Superior Parietal Lobule | 7 |
|  | 36 -36 38 | Inferior Parietal Lobule | 40 |
|  | 48 -68 10 | Middle Occipital Gyrus | 19 |
| 9568 | 27 5 53 | Superior Frontal Gyrus | 6 |
|  | 40 13 45 | Middle Frontal Gyrus | 8 |
| 6152 | -29 2 60 | Middle Frontal Gyrus | 6 |
|  | -22 -12 46 | Superior Frontal Gyrus | 6 |
|  | -26 -14 46 | Dorsal Precentral Gyrus | 6 |
|  | -34 -8 50 | Dorsal Precentral Gyrus | 6 |
| 3336 | -47 -67 -5 | Inferior Occipital Gyrus | 19 |
|  | -44 -72 8 | Middle Occipital Gyrus | 19 |
| 3128 | -48 14 27 | Inferior Frontal Gyrus | 6 |
|  | -42 24 30 | Middle Frontal Gyrus | 9 |
|  | -44 -2 32 | Precentral Gyrus | 6 |
| 1520 | 46 9 26 | Inferior Frontal Gyrus | 9 |
| 976 | -2 14 48 | SMA | 6 |
| 736 | -32 20 -14 | Inferior Frontal Gyrus/Insula | 47/13 |

### Activations related to time processing

#### Time-related activations

The meta-analysis of the studies on timing processing included 1262 foci from 114 experiments for a total of 1703 participants (see Table 3, Figure 3a). The minimum cluster size for the cluster to be considered statistically significant was 1120 mm3. Two quasi-symmetrical big clusters, one in the right hemisphere (22192 voxels) and one in the left hemisphere (15320 voxels), revealed a consistent strong activation of the basal ganglia: globus pallidum, putamen extending to the right caudate nucleus. Bilateral activations of thalamus, anterior insula and inferior frontal gyri were also detected within the same clusters. Medially, SMA regions (Pre-SMA and SMA-proper) were consistently activated (cluster of 15120 voxels). On the lateral brain surface, areas of high convergence among timing studies were found in the inferior parietal gyrus of both hemispheres (4296 voxels and 2768 voxels in the right and left hemispheres, respectively) including the intraparietal sulci (bilaterally, but larger in the right hemisphere). Furthermore, the precentral gyri were activated (3472 voxels in the left hemisphere and 2768 in the right one). Additional clusters of activation were found in the right superior temporal regions (1232 voxels), and in the right middle frontal gyrus (1192 voxels). Areas of overlap were detected in right and left cerebellar hemispheres.

**Table 3.** Significant activation likelihood clusters for the analysis of time processing.

| Cluster | Coordinates | Brain Region | Broadman Area |
| --- | --- | --- | --- |
| 22192 | 16 8 -2 | Pallidum | 51 |
|  | 54 12 16 | Inferior Frontal Gyrus | 44 |
|  | 22 6 6 | Putamen | 49 |
|  | 52 14 12 | Inferior Frontal Gyrus | 44 |
|  | 34 22 -2 | Insula | 13 |
|  | 12 -14 8 | Thalamus | 50 |
|  | 48 14 -4 | Insula | 13 |
|  | 56 14 -6 | Superior Temporal Pole | 22 |
|  | 10 -20 -6 | Red Nucleus | -- |
| 15320 | -14 -18 4 | Thalamus | 50 |
|  | -30 24 0 | Insula | 13 |
|  | -52 10 4 | Inferior Frontal Gyrus | 44 |
|  | -22 4 6 | Putamen | 49 |
|  | -18 8 -6 | Pallidum | 51 |
|  | -26 -8 -4 | Putamen | 49 |
|  | -28 -14 4 | Putamen | 49 |
| 15120 | 0 6 56 | SMA | 6 |
|  | 4 18 48 | SMA | 8 |
|  | 10 16 36 | SMA | 8 |
| 4296 | 44 -44 44 | Inferior Parietal Lobule | 40 |
|  | 54 -36 48 | Inferior Parietal Lobule | 40 |
| 4296 | 32 -62 -28 | Cerebellum | --- |
|  | 22 -56 -26 | Cerebellum | --- |
|  | 42 -58 -34 | Cerebellum | --- |
| 3472 | -48 -4 50 | Precentral Gyrus | 6 |
| 2928 | -26 -62 -26 | Cerebellum | --- |
| 2768 | -36 -22 54 | Precentral Gyrus | 4 |
|  | -50 -24 54 | Postcentral Gyrus | 1 |
|  | -44 -34 46 | Inferior Parietal Lobule | 40 |
|  | -36 -46 40 | Inferior Parietal Lobule | 40 |
| 1232 | 66 -36 16 | Superior Temporal Gyrus | 22 |
|  | 58 -34 8 | Superior Temporal Gyrus | 22 |
| 1192 | 40 38 28 | Middle Frontal Gyrus | 9 |

**Figure 3.**
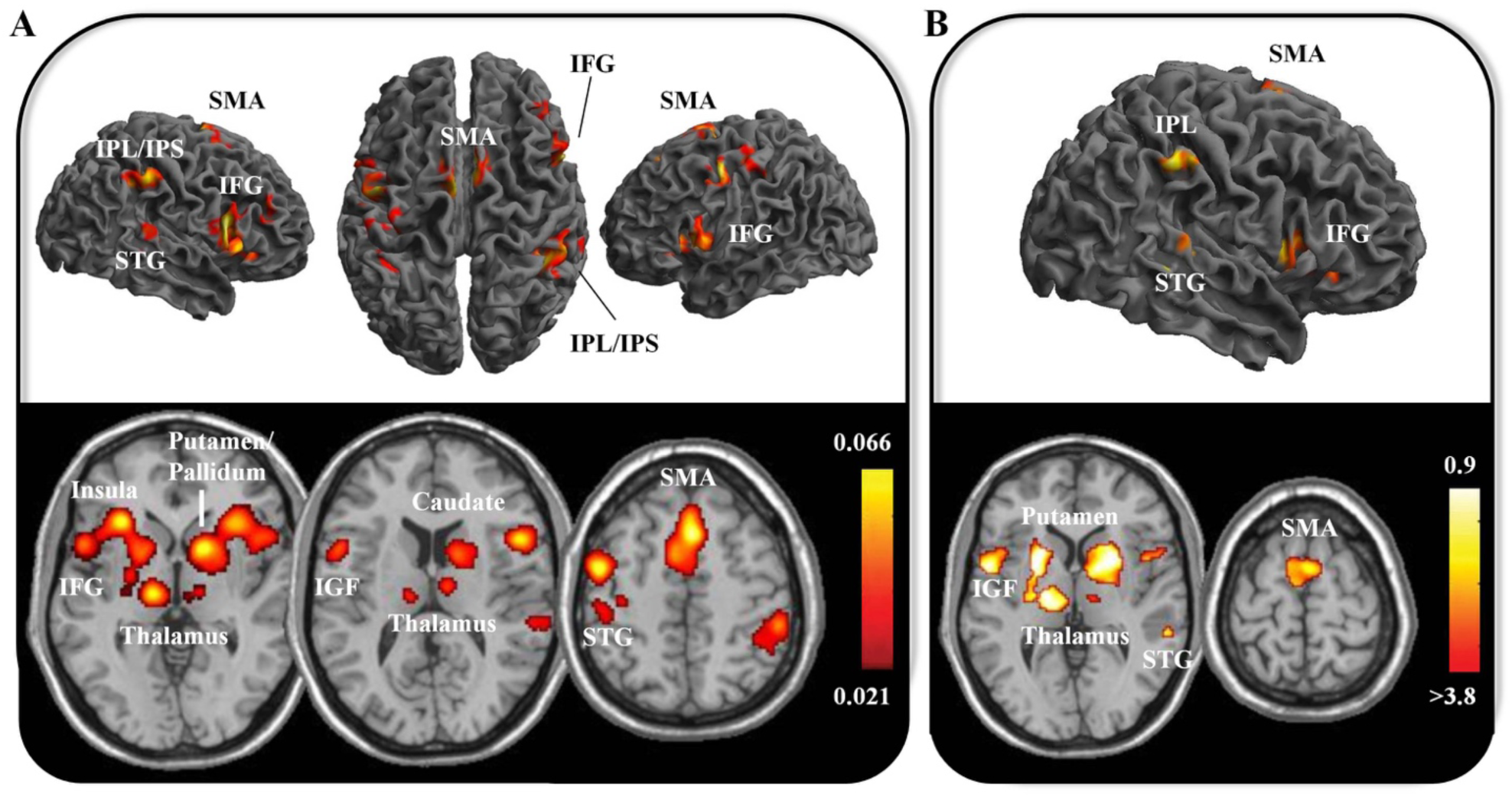
Time related brain activations. A) Brain regions that are activated during tasks requiring time processing. Colors indicate the ALE values for each voxel above the threshold (where yellow indicates the most significant ALE values); B) Brain regions where the convergence of coordinates of activation is higher during tasks requiring processing of time rather than space (Time > Space). To simplify the interpretation of ALE contrast images, they are converted to z scores to show their significance instead of a direct ALE subtraction (z-score for each voxel above the threshold). IPL = Inferior Parietal Lobule; IPS = Intra Parietal Sulcus; STG= Superior Temporal Gyrus; SMA = Supplementary Motor Area; IFG = Inferior Frontal Gyrus.

#### Time minus space related activations

A meta-analysis that identified brain activations that were activated to a greater extent for time than for space (Time > Space direct contrast) was then run. This meta-analysis identified a set of areas that were specific for time processing (Table 4, Figure 3b). These included all the basal ganglia structures identified in the previous meta-analysis of timing. In particular, a big cluster (7344 voxels) on the left hemisphere involved the pallidum, putamen and thalamus. In the right hemisphere a big cluster (7848 voxels) involved the pallidum and putamen, extending to the caudate nucleus. The thalamus is involved in a separate small cluster (232 voxels). The cerebellum regions still showed consistent activations bilaterally (4152 voxels and 2368 voxels in the right and left hemisphere, respectively). Notably, no activation over insular cortices were detected.

**Table 4.**
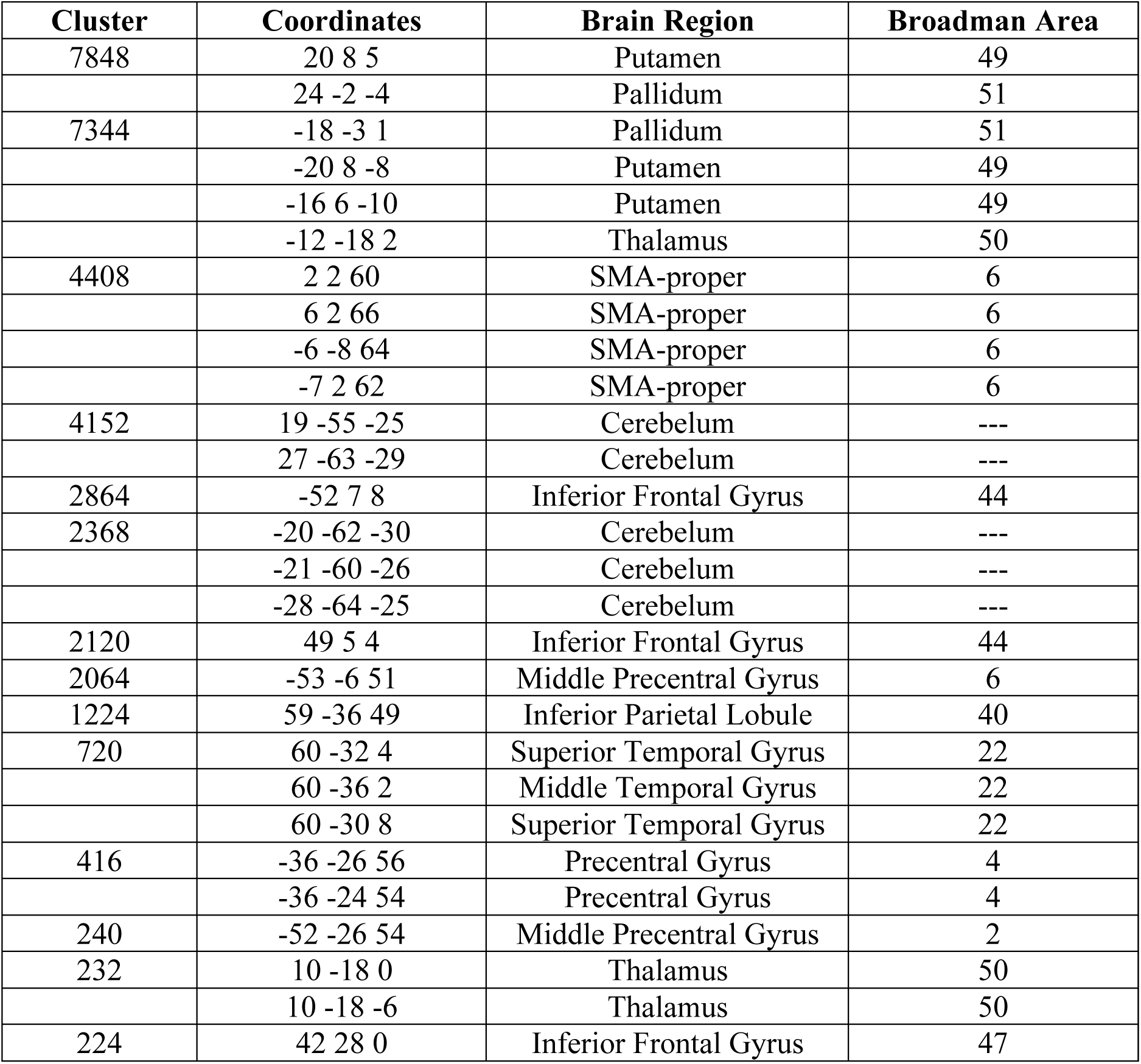
Significant activation likelihood clusters for the time > space analysis.

On the cortical medial surface, areas of convergence on SMA-proper (a big cluster of 4408 voxels) were observed, whereas laterally, we found consistent activations in the bilateral frontal opercula (BA 44) and left inferior parietal lobule (BA 40). The other, smaller, clusters of activation in the right middle and superior temporal regions, and in the left pre-central gyrus were still detected in this meta-analysis.

### Common activations and gradients

The conjunction analysis isolated those areas that were commonly activated in space and time studies (see Table 5, Figure 4). The analysis showed strong areas of convergence bilaterally in anterior insular cortices (clusters of 2480 and 2320 voxels for right and left hemisphere, respectively). Also, areas of convergence were identified bilaterally in: 1) the SMA regions, most prominently the pre-SMA (7904 voxels), 2) in the right frontal operculum (2200 voxels), and 3) bilaterally in a parietal region centered around the intra-parietal sulci (1776 and 856 voxels, for right and left hemisphere, respectively); 4) the left precentral gyrus (336 voxels, BA 6).

**Table 5.** Significant activation likelihood clusters for conjunction space & time analysis.

| Cluster | Coordinates | Brain Region | Broadman Area |
| --- | --- | --- | --- |
| 7904 | 4 18 48 | Pre-SMA | 8 |
|  | 0 8 54 | Pre-SMA | 6 |
| 2480 | 36 22 0 | Insula | 13 |
| 2320 | -30 24 -2 | Insula | 13 |
| 2200 | 54 12 20 | Inferior Frontal Gyrus | 44 |
| 1776 | 42 -44 44 | Inferior Parietal Lobule | 40 |
| 856 | -44 -34 46 | Postcentral Gyrus | 40 |
|  | -36 -46 40 | Inferior Parietal Lobule | 40 |
| 336 | -42 0 50 | Precentral Gyrus | 6 |

**Figure 4.**
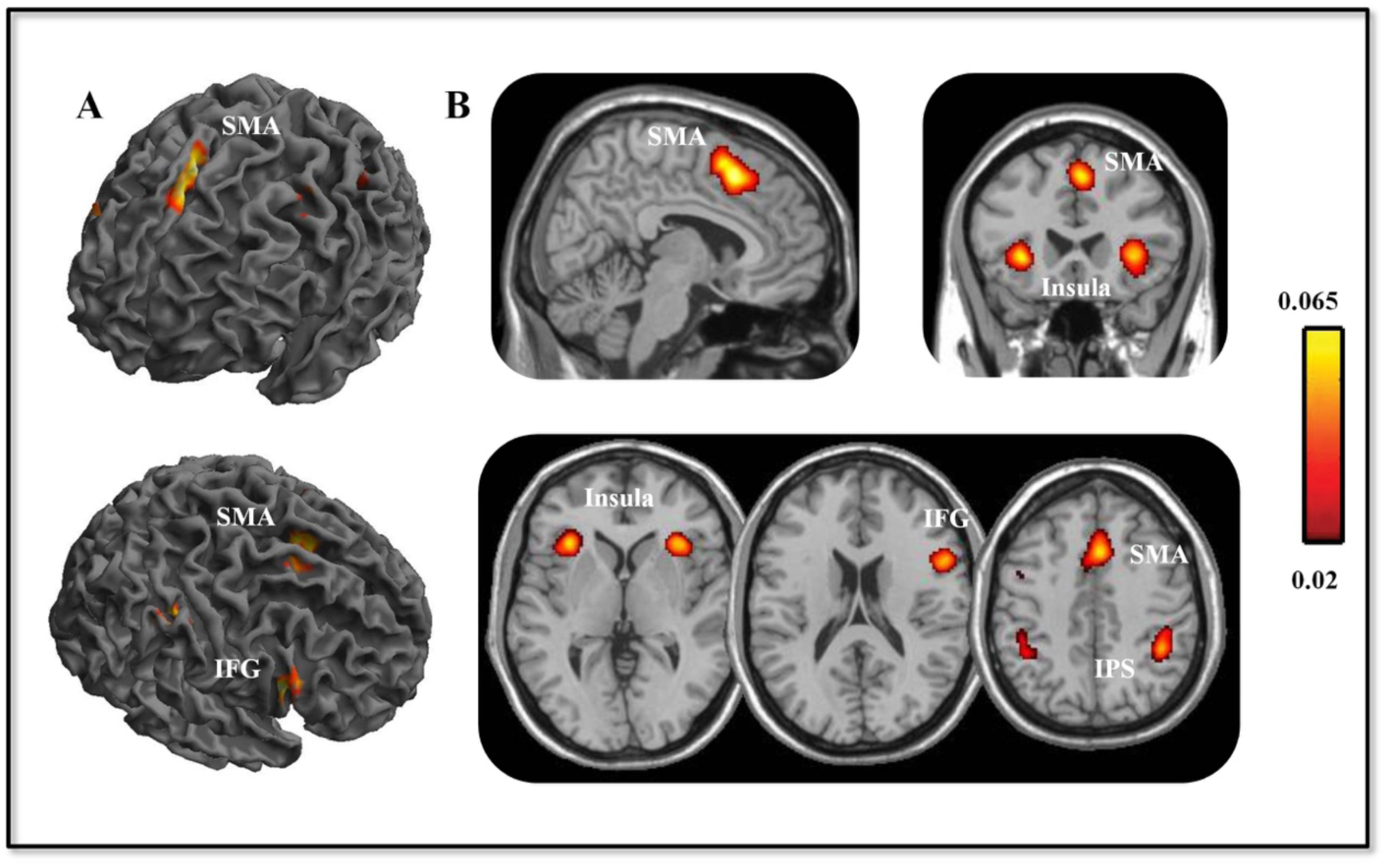
Conjunction analysis. This figure illustrates common ALE activations for both space and time. A) results displayed on 3D renders; B) results displayed on 2D sections and in particular on sagittal and axial section in the upper line and on the axial section on the lower line. Colors indicated the ALE values for each voxel above the threshold (where yellow indicates the most significant ALE values). SMA = Supplementary Motor Area; IFG = Inferior Frontal Gyrus; IPS = Intra Parietal Sulcus.

Notably, as illustrated in Figure 5, all the regions of common activation but insula (i.e., SMA, right frontal operculum, IPS, and left precentral gyrus) represent the “intersection” of topographical gradients, along which space and time are mapped and organized in the brain.

**Figure 5.**
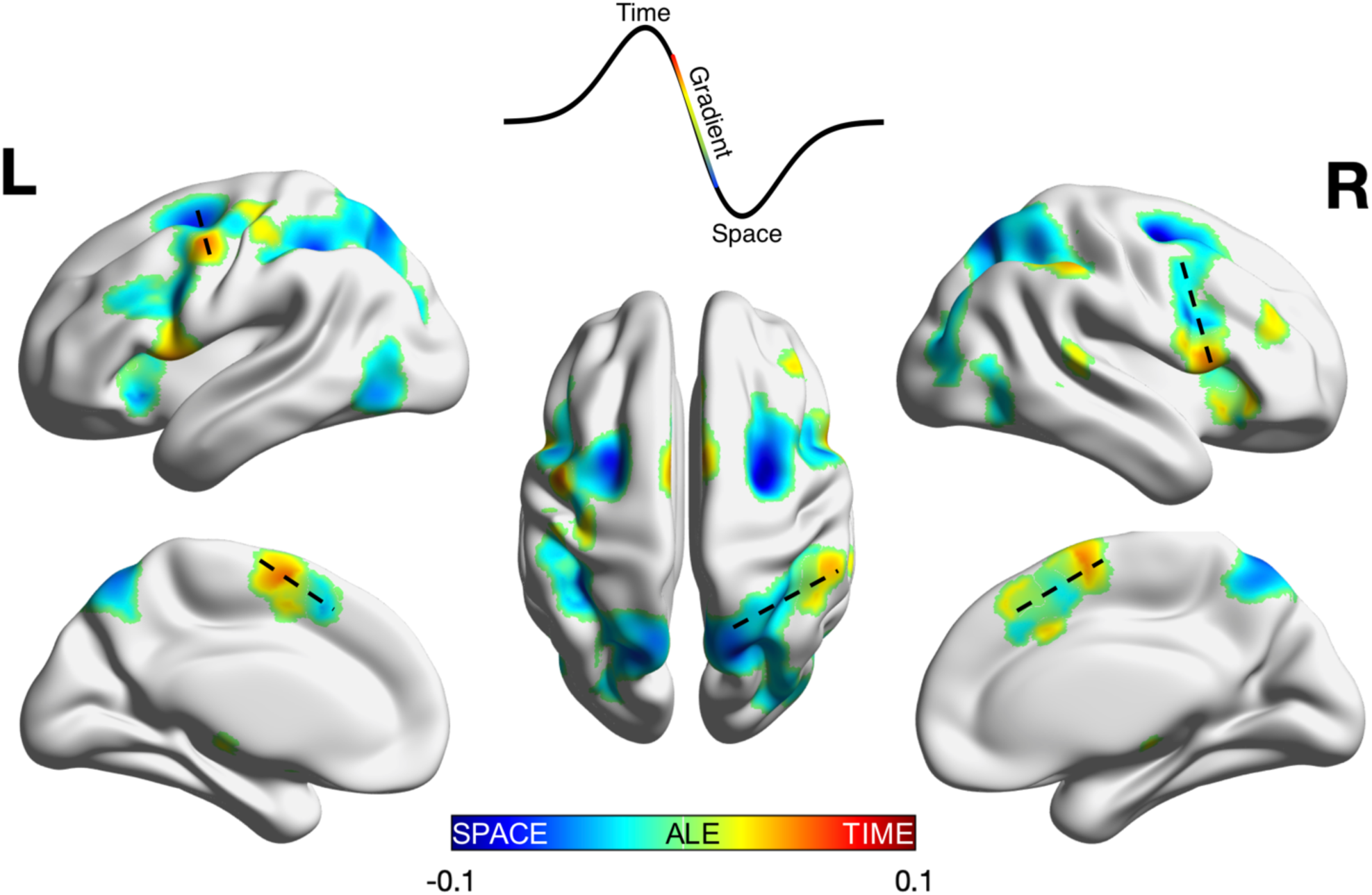
Gradient Analysis. Surface visualization of the overlap between Space and Time meta-analyses. ALE values for Space were set to negative numbers and added to ALE values from the Time meta-analysis, effectively subtracting one from the other. Gradients were identified as transition zones from positive-to-negative values, as indicated by dashed lines. Altogether, gradients were observed within the bilateral SMA, right inferior-to-superior parietal cortex, right inferior-to-superior prefrontal cortex, and the left precentral gyrus.

The SMA regions show an anterior-posterior topographical gradient, with space activating more anterior regions (i.e., pre-SMA) and time activating more posterior regions (thus, SMA-proper to greater extent). Frontal and parietal regions show a dorsal-ventral gradient. Indeed, space is associated with activation of dorsal frontal and parietal regions, whereas time is more likely to activate ventral frontal and parietal regions. In particular, on the right hemisphere, space activates more the precuneus and superior parietal lobule, whereas time activates more the left inferior parietal lobule. Both space and time activate the IPS, which is situated right in the middle and separates inferior from superior lobule. Furthermore, space is mainly supported by right superior and middle frontal area while time is mainly supported by right inferior frontal area. The frontal operculum is commonly activated by both the domains. Finally, on the left hemisphere, dorsal regions of the left precentral gyrus (i.e., dorsal premotor cortex, PMd) are activated for space-related processes whereas ventral regions of the precentral gyrus are activated for time-related processes. Notably, no gradients were observed in regions that did not contain areas of common activation between space and time.

## Discussion

According to the ATOM, formulated by Walsh more than fifteen years ago, there is a common system of magnitude in the brain that comprises regions – such as parietal cortex – shared by space, time and other magnitudes (Beudel, Renken, Leenders, & de Jong, 2009; Bueti & Walsh, 2009; Walsh, 2003). The present meta-analysis clearly identified the presence of a set of brain regions that are commonly recruited in both space and time. This system includes bilateral insula, the pre-SMA, the right frontal operculum and the intraparietal sulci. Our study supports and updates the ATOM theory, as it showed not only overlapping activations between space and time but also revealed that spatial and temporal processing is arranged and organized along well-defined spatial gradients in the brain (see Figure 5). For this reason, we now refer to Walsh’s theory as ‘*GradiATOM*’ (Gradient Theory of Magnitude).

### ‘GradiATOM’: Functional gradients underlying space and time

We found that pre-SMA, right inferior frontal gyrus (IFG), left precentral gyrus and intraparietal sulci represent the areas of activation overlap with spatial gradients, along which space and time are mapped and organized in the brain. More specifically, the SMA showed an anterior-posterior gradient, with space activating more anterior regions (i.e., pre-SMA) and time activating more posterior regions (thus, SMA-proper to greater extent). Frontal and parietal regions showed a dorsal-ventral gradient. Space processing is supported by dorsal frontal and parietal regions, whereas time is more likely to recruit ventral frontal and parietal regions.

A recent study by Eickhoff et al. (2018) emphasized that the brain is characterized by multiple topographies at different scales, ranging from local properties of brain structures to large-scale networks. Gradients consist of a functional and/or anatomical segregation into distinct subdivisions, which emerges through local differentiation. Functionally, a large body of evidence demonstrated a gradient principle along the anterior-posterior axis of the frontal lobes on the basis of the abstractness of action representations (Badre & D’Esposito, 2009). Indeed, neurons of more-anterior prefrontal cortex (PFC) encodes abstract action goals, while neurons of more-posterior regions encode more-concrete details of action (Badre & D’Esposito, 2009). Moreover, more-anterior regions interact with more-posterior regions hierarchically, with anterior regions being more likely to influence posterior regions than *vice versa* (Badre, Hoffman, Cooney, & D’Esposito, 2009; Fuster, 2004). Spatial gradients related to concreteness-abstractness are not confined to the PFC. There is now consistent evidence for global hierarchical gradients that separates concrete (perceptual and physical) categories in sensorimotor regions from more abstract concepts encoded in transmodal areas (Huth et al., 2016; Huntenburg et al., 2018). Hierarchical gradients emerge from unimodal sensorimotor areas and radiates toward transmodal, higher-order areas, culminating in the default mode network and salience network (Huntenburg et al., 2018; Margulies et al., 2016; Vázquez-Rodríguez et al., 2019). Along these gradients, low-level sensory features are increasingly abstracted and integrated with concepts from other systems.

In our study, we found functional gradients in a set of frontal and parietal regions. Importantly, such gradients contain areas of common activations, which represent the ‘areas of conjunction’ of space-related and time-related activations. These areas would thus play a similar role in both space and time domain, whereas sub-areas in the gradients would be specialized to process and act efficiently and selectively on space and time material. Indeed, previous neural recording studies in non-human primates have revealed subpopulations of neurons tuned specifically for duration or spatial distance, as well neurons tuned for both dimensions, all within the same cortical region (Genovesio, Seitz, Tsujimoto, & Wise, 2016; Genovesio, Tsujimoto, & Wise, 2012).

Following this logic and on the basis of previous studies, fronto-parietal networks would have a domain-general role, linked to allocating attention toward spatial and temporal stimuli and their representation (Corbetta, Patel, & Shulman, 2008; Coull & Nobre, 2008; Coull & Nobre, 1998; Luckmann et al., 2014). By contrast, sub-regions of frontal and parietal regions would be preferentially activated by spatial (dorsal fronto-parietal regions) and temporal (ventral fronto-parietal regions) information in order to represent it (e.g., Hagler & Sereno, 2006). Topographic maps or ‘prioritized maps of space’ were indeed discovered in frontal and parietal regions, especially in dorsal regions such as frontal eye fields and superior parietal areas close to intraparietal sulcus (Hagler & Sereno, 2006; Serences & Yantis, 2007; Szczepanski et al., 2010). For example, a study showed that, similar to visual cortex, human fronto-parietal cortices contain topographic representations of eccentricity and polar angle, which are organized into clusters so that to represent all the gradients of polar angle of the contralateral visual field (Mackey, Winawer, & Curtis, 2017). Likewise, very recent evidence identified topographic timing maps or chronotopic maps in several brain regions, including not only sensory cortices but also frontal, parietal areas and SMA (Harvey, Damoulin, Fracasso, & Paul, 2019; Protopapa et al., 2019).

The SMA showed an anterior-posterior gradient, with more-anterior regions (i.e., pre-SMA) supporting space and more-posterior regions (thus, SMA-proper to greater extent) supporting time. Regions in the central part of SMA were instead found commonly activated by both space and time. SMA has been shown to mediate the sequential integration of elements into hierarchically organized-representations across a variety of stimuli (temporal, spatial, linguistic, numerical etc.; Cona & Semenza, 2017). Therefore, SMA might act to sequentially integrate information in both space and time domains, with subregions of SMA being active preferentially for spatial elements (i.e., the anterior regions) and temporal elements (i.e., the posterior regions) (see paragraph below for a more detailed discussion).

Finally, a smaller cluster of common activations was also found over the left dorsal premotor (PMd) region, along which a dorsal-ventral gradient occurs. Dorsal areas of PMd were more activated for spatial tasks, whereas ventral areas of PMd were more activated for temporal tasks. PMd is a heterogenous functional region that was shown to be composed of five different modules supporting a variety of cognitive and motor functions (Genon et al., 2018). An anterior-posterior organization of the PMd modules was found to reflect a cognitive-motor gradient. Further, the central PMd module was associated with functions as mental rotation, visuo-spatial attention and spatial cognition, whereas the ventral module mediates functions that rely more upon time, as music cognition and language (Genon et al., 2018). Our findings support the mosaic nature of this region, showing that the dorsal PMd module is mainly devoted to spatial material, while the ventral PMd module is associated with temporal material.

Taken together, these results brought the first evidence for an organization along gradients for space-related and time-related neural processes. This spatial proximity would ensure the interplay and integration of space and time information into a coherent representation of the external world that will be used for preparing the appropriate action. In such a way the *GradiATOM* theory nicely fits with the original ATOM view that “space and time are coupled metrics for action and it would be very surprising if they were not in close proximity in the brain” (p. 1832, Bueti & Walsh, 2009). Furthermore, our theory updates the ATOM view, showing that space and time representations are distributed in the brain along an anterior-posterior axis in the SMA regions, and a dorsal-ventral axis in the fronto-parietal regions. Based on the evidence for an anterior-posterior and dorsal-ventral processing hierarchy in the PFC (Badre et al., 2009; Schumacher et al., 2019), it is plausible to assume a hierarchy in space-time processing as well, where temporal inputs are processed and manipulated “*in ministerium*” of forming/enriching spatial representations to greater extent than *vice versa*. Gradient organization thus facilitates hierarchical transformations by enabling space- and time-related neural populations to interact with each other over minimal distances (Harvey & Dumoulin, 2017).

### Common neural network for space and time

As described above, we identified five brain regions that were commonly active across space and time conditions: anterior insular cortices, the right IFG, bilateral SMA, the left precentral gyrus and IPS. Since these regions are evident in both the two domains, we suggest they play a pivotal, domain-general, role in processes that act on time and space material (e.g., fronto-parietal network in attention). The specificity of the material to process (e.g., attention to maps of space or attention to time) would be instead reflected in the gradient, where distinct regions are selectively more activated for spatial or temporal elements. All these regions but insula, indeed, fall within the gradients and represent the point of conjunction between space-related activations and time-related activations.

The anterior insula and right frontal operculum are part of the ‘salience network’ (Seeley et al., 2007) and ‘cingulo-opercular control’ network (Dosenbach et al., 2008). More specifically, the anterior insula is involved in the transient identification of salient and/or relevant (either internal or external) stimuli in order to guide thoughts and behaviour (Seeley et al., 2007). Activity in frontal operculum has been instead associated with updating and prioritizing processes (Myers, Stokes, & Nobre, 2017; Visalli, Capizzi, Ambrosini, Mazzonetto, & Vallesi, 2019). In particular, an elegant study carried out by Visalli and collaborators (2019) was able to decouple the updating and surprise components of temporal expectations and to disentangle their related neural substrates. They found that the updating component was associated mainly with fronto-parietal regions, such as inferior frontal gyrus, whereas surprise component modulated areas of the cingulo-opercular control network, such as anterior insula. This recent finding supports the interpretation given in our schematic model of spatial processing (Cona & Scarpazza, 2019), according to which representations of space are prioritized by frontal operculum and the insula on the basis of the relevance of the individuals’ goals and the salience of the external stimuli. As these regions are shared between the two domains, and based on the recent evidence in the literature (Dosenbach et al., 2008; Myers et al., 2017; Seeley et al., 2007; Visalli et al., 2019), such interpretation can be extended to time processes. The ‘prioritizing’ operations can indeed be meant as the process by which any information representation (either spatial or temporal) is updated by frontal operculum based on the saliency inputs sent by insular cortices. Notably, the insula was the one region of conjunction that did *not* exhibit a gradient representation. This finding may indicate that the insula represents a domain-general prioritizing process in both domains.

Overlapping activations between space and time domains were also shown over SMA regions. This finding is in line with the view of a domain-general role of SMA regions in sequencing operations (Cona & Semenza, 2017). SMA is involved in integrating sequential elements into hierarchically organized-representations regardless of the nature of such elements (temporal, spatial, linguistic, numerical etc.; Cona & Semenza, 2017). In the space domain, previous studies found indeed that SMA involvement (and mainly pre-SMA) strictly depends on the complexity of visuo-spatial sequential transformations, such as those implied in mental rotation (e.g., (Cona et al., 2017; Milivojevic et al., 2009; Richter et al., 2000). Likewise, in time domain, build-up activity over SMA was found to be influenced by temporal duration (Bendixen, Grimm, & Schroger, 2005; Macar & Vidal, 2002; Macar, Vidal, & Casini, 1999). This pattern of activity has led several researchers to indicate the SMA as the temporal accumulator (Casini & Vidal, 2011; Coull et al., 2015). Interestingly, recent evidence revealed a clear gradient organization along the anterior-posterior axis of SMA, with anterior and posterior SMA sub-regions showing preferential activations for short and long durations, respectively (Protopapa et al., 2019; see also Harvey et al., 2019). This duration-sensitive tuning organized in a gradient might be a good candidate to sequentially integrate temporal pulses into a representation of durations.

Notably, another functional gradient was observed over the SMA, wherein motor timing tasks were associated with the SMA-proper activation, while perceptual timing tasks tended to activate pre-SMA (Schwartze et al., 2012; Wiener et al., 2011; Wiener et al., 2010). This gradient would also partially explain the functional gradient we observed in SMA regions as a function of space-time processing. Indeed, temporal tasks are more likely to include a motor component (e.g., paced finger tapping, interval production/reproduction tasks etc.) as compared with spatial tasks. Even if it is true that the control conditions in timing studies included similar motor tasks, nonetheless collateral motor-related operations (such as, programming rhythmic sequences of actions) might be not completely removed in the subtraction analysis.

The intraparietal sulci showed a consistent overlap of activation between space and time. This is probably the first brain region that has received interest for its role in representing magnitude related to space, time, number and other magnitudes (Bueti & Walsh, 2009; Walsh, 2003; Weger & Pratt, 2008). As mentioned before, intraparietal sulci are the areas of overlaps between the more-dorsal parietal activations, related to space, and more-ventral parietal activations related to time. There is a large consensus that intraparietal sulci support general representation mechanisms in both temporal domain and spatial domain, as they contain both topographic maps of space and chronotopic maps (Hagler & Sereno, 2006; Hayashi et al., 2015; Jerde & Curtis, 2013; Mackey et al., 2017; Teghil et al., 2019). The coexistence of both types of map in the same brain region makes the intraparietal sulci the ideal candidate for operations like transformation and integration of spatio-temporal information.

### Summary and conclusion

The present study provides the first meta-analysis of neuroimaging studies on space and time processing in order to unveil possible overlapping activations between the two domains. We found a set of brain regions that were consistently and commonly activated in space and time, and that comprised bilateral insula, the pre-SMA, the right frontal operculum, the left precentral gyrus and the intraparietal sulci. These regions might be the best candidates to form the ‘core’ magnitude circuit. Importantly, the activations in these regions but the insula represent the overlaps of patterns of magnitude-related activations that are distributed as gradients, along which space and time were organized in the brain. Frontal and parietal regions showed a dorsal-ventral gradient, with space processing being supported by dorsal frontal and parietal regions, and time being associated more likely to ventral frontal and parietal regions. An anterior-posterior gradient was observed over SMA regions, with space activating more-anterior regions (i.e., pre-SMA) and time activating more-posterior regions (thus, SMA-proper to greater extent). We suggest that the brain regions shared by space and time might play a similar process in the two domains, while the gradients observed along such regions would reflect the specific material to process (i.e., spatial versus temporal information).

A limitation of the present study is that the gradient organization has been found at the group level. Therefore, further studies are needed to better characterize these topographic neural gradients possibly using a within-subject design. Nonetheless, our study is important as it provides the first clues that time and space processes are likely to be organized along gradients. We hypothesized that the spatial proximity derived from such gradient organization would facilitate the integrations of magnitude representations by enabling space- and time-related neural populations to interact with each other.

## Acknowledgements

We thank Noemi Terruzzi for her help in extracting the data. “This work was carried out within the scope of the project “use-inspired basic research”, for which the Department of General Psychology of the University of Padova has been recognized as “Dipartimento di eccellenza” by the Ministry of University and Research”.

