## Supplementary file A for "From *ATOM* to *GradiATOM*: Cortical gradients support time and space processing as revealed by a meta-analysis of neuroimaging studies"

**Supplementary Information A**

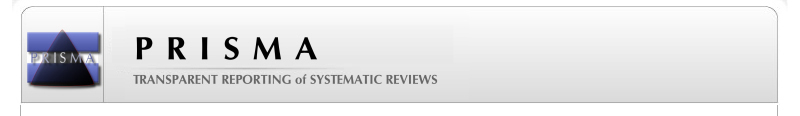
**PRISMA 2009 Flow Diagram**

**(last update March 2018)**

**Screening**

**Included**

**Eligibility**

**Identification**

Additional records identified through other sources (“related articles” function in Pubmed, or articled included within the references
(n = 167)

Records identified through database searching
(n = 290)

Records after duplicates removed
(n = 457)

Records screened
(n = 457)

Records excluded
(n = 181)

Full-text articles assessed for eligibility
(n= 276)

Studies included in qualitative synthesis
(n = 133)

Studies included in quantitative synthesis (meta-analysis)
(n =110)

Full-text articles excluded

(n=11: no fMRI;

n=3: pathological population only;

n=15: no coordinates reported;

n=9: functional connectivity o machine learning analysis;

n=4: task mixing time and space;

n=7: task does not involve space;

n=42: the isolated contrast does not involve space;

n=23: LTM&navigation;

n=8: small sample size;

n=41: ROI studies;

n=2: deactivation only reported;

n=1: negative results)

(n = 166)
