## Supplementary file B for "From *ATOM* to *GradiATOM*: Cortical gradients support time and space processing as revealed by a meta-analysis of neuroimaging studies"

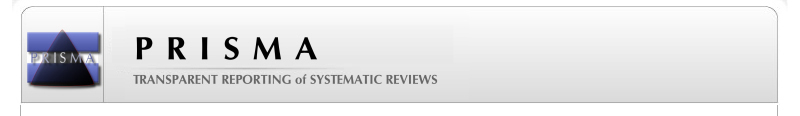
**Supplementary Information B**

**PRISMA 2009 Flow Diagram**

**(last update February 2019)**

**Screening**

**Included**

**Eligibility**

**Identification**

Full-text articles excluded, with reasons:

(n=14: no fMRI;

n=4: pathological population only;

n=7: no coordinates reported;

n=2: functional connectivity o machine learning analysis;

n=3: task mixing time and space;

n=15: task does not involve time;

n=17: the isolated contrast does not involve time;

n=20: ROI studies;

reported)

(n = 82)

Records identified through database searching
(n = 595)

Additional records identified through other sources (cross reference; “similar articles” function)
(n = 27)

Records after duplicates removed
(n = 622)

Records screened
(n = 622)

Records excluded
(n = 430)

Full-text articles assessed for eligibility
(n = 192)

Studies included in qualitative synthesis
(n = 110)

Studies included in quantitative synthesis (meta-analysis)
(n = 110)
