## Supplementary file C for "From *ATOM* to *GradiATOM*: Cortical gradients support time and space processing as revealed by a meta-analysis of neuroimaging studies"

**Supplementary Information C**

**List of Included Studies for the “Space” meta-analysis**

(Alivisatos & Petrides, 1997; Arrington, Carr, Mayer, & Rao, 2000; Bahlmann, Schubotz, Mueller, Koester, & Friederici, 2009; Barnes et al., 2000; Beauchamp, Petit, Ellmore, Ingeholm, & Haxby, 2001; Belger et al., 1998; Bien & Sack, 2014; Bonda, Petrides, Frey, & Evans, 1995; Braga, Fu, Seemungal, Wise, & Leech, 2016; Bushara et al., 1999; Carpenter, Just, Keller, Eddy, & Thulborn, 1999; Chen, Weidner, Vossel, Weiss, & Fink, 2012; Cohen et al., 1996; Committeri et al., 2004; Corbetta et al., 1998; Corbetta, Miezin, Shulman, & Petersen, 1993; Coull & Nobre, 1998; Courtney, Ungerleider, Keil, & Haxby, 1996; Curtis, Rao, & D'Esposito, 2004; D'Esposito et al., 1998; Doricchi, Macci, Silvetti, & Macaluso, 2010; Dupont et al., 1998; Ecker, Brammer, David, & Williams, 2006; Egner et al., 2008; Fagioli & Macaluso, 2009, 2016; Fairhall, Indovina, Driver, & Macaluso, 2009; Fusser et al., 2011; Galati, Committeri, Sanes, & Pizzamiglio, 2001; Galati et al., 2000; Geier, Garver, & Luna, 2007; Giesbrecht, Woldorff, Song, & Mangun, 2003; Gitelman et al., 1999; Halari et al., 2006; Harris et al., 2000; Hautzel et al., 2002; Humphreys & Lambon Ralph, 2017; Indovina & Macaluso, 2007; Johnston, Leek, Atherton, Thacker, & Jackson, 2004; Jonides et al., 1993; Jordan, Heinze, Lutz, Kanowski, & Jancke, 2001; Jordan, Wustenberg, Heinze, Peters, & Jancke, 2002; Kelley, Serences, Giesbrecht, & Yantis, 2008; Kim et al., 1999; Kincade, Abrams, Astafiev, Shulman, & Corbetta, 2005; S. M. Kosslyn, DiGirolamo, Thompson, & Alpert, 1998; S.M. Kosslyn, Thompson, Gitelman, & Alpert, 1998; Krumbholz, Nobis, Weatheritt, & Fink, 2009; LaBar, Gitelman, Parrish, & Mesulam, 1999; Lamm, Windischberger, Leodolter, Moser, & Bauer, 2001; Lamm, Windischberger, Moser, & Bauer, 2007; Lamp, Alexander, Laycock, Crewther, & Crewther, 2016; Leek, Yuen, & Johnston, 2016; Lepsien, Griffin, Devlin, & Nobre, 2005; Leung, Seelig, & Gore, 2004; Lipp et al., 2012; Macaluso, Eimer, Frith, & Driver, 2003; Macaluso, Frith, & Driver, 2002; Macaluso & Patria, 2007; Manoach et al., 2004; Milivojevic, Hamm, & Corballis, 2009; Nardo, Santangelo, & Macaluso, 2011; Natale, Marzi, & Macaluso, 2009; Nemmi, Boccia, Piccardi, Galati, & Guariglia, 2013; Ng et al., 2001; Nobre et al., 1997; Nystrom et al., 2000; Ogawa & Macaluso, 2015; Owen, Evans, & Petrides, 1996; Peelen, Heslenfeld, & Theeuwes, 2004; Podzebenko, Egan, & Watson, 2002, 2005; Ricciardi et al., 2006; Rosen et al., 1999; Rowe, Toni, Josephs, Frackowiak, & Passingham, 2000; Sala, Rama, & Courtney, 2003; Salmi, Rinne, Degerman, Salonen, & Alho, 2007; Santangelo, Olivetti Belardinelli, Spence, & Macaluso, 2009; Schmidt et al., 2007; Schubotz & von Cramon, 2001; Seurinck, Vingerhoets, de Lange, & Achten, 2004; Seurinck, Vingerhoets, Vandemaele, Deblaere, & Achten, 2005; Seydell-Greenwald, Ferrara, Chambers, Newport, & Landau, 2017; Shen, Hu, Yacoub, & Ugurbil, 1999; Simon et al., 2002; D. V. Smith et al., 2010; E. E. Smith, Jonides, & Koeppe, 1996; E. E. Smith et al., 1995; Summerfield, Lepsien, Gitelman, Mesulam, & Nobre, 2006; Tagaris et al., 1997; Tagaris et al., 1998; Thiel, Zilles, & Fink, 2004; Thomason et al., 2009; Thomsen et al., 2000; Umla-Runge, Zimmer, Krick, & Reith, 2011; Vallar et al., 1999; Vandenberghe, Gitelman, Parrish, & Mesulam, 2001; Vanrie, Beatse, Wagemans, Sunaert, & Van Hecke, 2002; Ventre-Dominey et al., 2005; Vingerhoets, de Lange, Vandemaele, Deblaere, & Achten, 2002; Walter et al., 2003; Weeks et al., 1999; Weiss et al., 2003; Windischberger, Lamm, Bauer, & Moser, 2003; Wraga, Thompson, Alpert, & Kosslyn, 2003; Yantis et al., 2002; Zacks, Rypma, Gabrieli, Tversky, & Glover, 1999; Zaehle et al., 2007)

Alivisatos, B., & Petrides, M. (1997). Functional activation of the human brain during mental rotation. *Neuropsychologia, 35*(2), 111-118. doi:10.1016/s0028-3932(96)00083-8

Arrington, C. M., Carr, T. H., Mayer, A. R., & Rao, S. M. (2000). Neural mechanisms of visual attention: object-based selection of a region in space. *J Cogn Neurosci, 12 Suppl 2*, 106-117. doi:10.1162/089892900563975

Bahlmann, J., Schubotz, R. I., Mueller, J. L., Koester, D., & Friederici, A. D. (2009). Neural circuits of hierarchical visuo-spatial sequence processing. *Brain Res, 1298*, 161-170. doi:10.1016/j.brainres.2009.08.017

Barnes, J., Howard, R. J., Senior, C., Brammer, M., Bullmore, E. T., Simmons, A., . . . David, A. S. (2000). Cortical activity during rotational and linear transformations. *Neuropsychologia, 38*(8), 1148-1156. doi:10.1016/s0028-3932(00)00025-7

Beauchamp, M. S., Petit, L., Ellmore, T. M., Ingeholm, J., & Haxby, J. V. (2001). A parametric fMRI study of overt and covert shifts of visuospatial attention. *Neuroimage, 14*(2), 310-321. doi:10.1006/nimg.2001.0788

Belger, A., Puce, A., Krystal, J. H., Gore, J. C., Goldman-Rakic, P., & McCarthy, G. (1998). Dissociation of mnemonic and perceptual processes during spatial and nonspatial working memory using fMRI. *Hum Brain Mapp, 6*(1), 14-32.

Bien, N., & Sack, A. T. (2014). Dissecting hemisphere-specific contributions to visual spatial imagery using parametric brain mapping. *Neuroimage, 94*, 231-238. doi:10.1016/j.neuroimage.2014.03.006

Bonda, E., Petrides, M., Frey, S., & Evans, A. (1995). Neural correlates of mental transformations of the body-in-space. *Proc Natl Acad Sci U S A, 92*(24), 11180-11184. doi:10.1073/pnas.92.24.11180

Braga, R. M., Fu, R. Z., Seemungal, B. M., Wise, R. J., & Leech, R. (2016). Eye Movements during Auditory Attention Predict Individual Differences in Dorsal Attention Network Activity. *Front Hum Neurosci, 10*, 164. doi:10.3389/fnhum.2016.00164

Bushara, K. O., Weeks, R. A., Ishii, K., Catalan, M. J., Tian, B., Rauschecker, J. P., & Hallett, M. (1999). Modality-specific frontal and parietal areas for auditory and visual spatial localization in humans. *Nat Neurosci, 2*(8), 759-766. doi:10.1038/11239

Carpenter, P. A., Just, M. A., Keller, T. A., Eddy, W., & Thulborn, K. (1999). Graded functional activation in the visuospatial system with the amount of task demand. *J Cogn Neurosci, 11*(1), 9-24. doi:10.1162/089892999563210

Chen, Q., Weidner, R., Vossel, S., Weiss, P. H., & Fink, G. R. (2012). Neural mechanisms of attentional reorienting in three-dimensional space. *J Neurosci, 32*(39), 13352-13362. doi:10.1523/JNEUROSCI.1772-12.2012

Cohen, M. S., Kosslyn, S. M., Breiter, H. C., DiGirolamo, G. J., Thompson, W. L., Anderson, A. K., . . . Belliveau, J. W. (1996). Changes in cortical activity during mental rotation. A mapping study using functional MRI. *Brain, 119 ( Pt 1)*, 89-100. doi:10.1093/brain/119.1.89

Committeri, G., Galati, G., Paradis, A. L., Pizzamiglio, L., Berthoz, A., & LeBihan, D. (2004). Reference frames for spatial cognition: different brain areas are involved in viewer-, object-, and landmark-centered judgments about object location. *J Cogn Neurosci, 16*(9), 1517-1535. doi:10.1162/0898929042568550

Corbetta, M., Akbudak, E., Conturo, T. E., Snyder, A. Z., Ollinger, J. M., Drury, H. A., . . . Shulman, G. L. (1998). A common network of functional areas for attention and eye movements. *Neuron, 21*(4), 761-773. doi:10.1016/s0896-6273(00)80593-0

Corbetta, M., Miezin, F. M., Shulman, G. L., & Petersen, S. E. (1993). A PET study of visuospatial attention. *J Neurosci, 13*(3), 1202-1226.

Coull, J. T., & Nobre, A. C. (1998). Where and when to pay attention: the neural systems for directing attention to spatial locations and to time intervals as revealed by both PET and fMRI. *J Neurosci, 18*(18), 7426-7435.

Courtney, S. M., Ungerleider, L. G., Keil, K., & Haxby, J. V. (1996). Object and spatial visual working memory activate separate neural systems in human cortex. *Cereb Cortex, 6*(1), 39-49. doi:10.1093/cercor/6.1.39

Curtis, C. E., Rao, V. Y., & D'Esposito, M. (2004). Maintenance of spatial and motor codes during oculomotor delayed response tasks. *J Neurosci, 24*(16), 3944-3952. doi:10.1523/JNEUROSCI.5640-03.2004

D'Esposito, M., Aguirre, G. K., Zarahn, E., Ballard, D., Shin, R. K., & Lease, J. (1998). Functional MRI studies of spatial and nonspatial working memory. *Brain Res Cogn Brain Res, 7*(1), 1-13. doi:10.1016/s0926-6410(98)00004-4

Doricchi, F., Macci, E., Silvetti, M., & Macaluso, E. (2010). Neural correlates of the spatial and expectancy components of endogenous and stimulus-driven orienting of attention in the Posner task. *Cereb Cortex, 20*(7), 1574-1585. doi:10.1093/cercor/bhp215

Dupont, P., Vogels, R., Vandenberghe, R., Rosier, A., Cornette, L., Bormans, G., . . . Orban, G. A. (1998). Regions in the human brain activated by simultaneous orientation discrimination: a study with positron emission tomography. *Eur J Neurosci, 10*(12), 3689-3699. doi:10.1046/j.1460-9568.1998.00376.x

Ecker, C., Brammer, M. J., David, A. S., & Williams, S. C. (2006). Time-resolved fMRI of mental rotation revisited--dissociating visual perception from mental rotation in female subjects. *Neuroimage, 32*(1), 432-444. doi:10.1016/j.neuroimage.2006.03.031

Egner, T., Monti, J. M., Trittschuh, E. H., Wieneke, C. A., Hirsch, J., & Mesulam, M. M. (2008). Neural integration of top-down spatial and feature-based information in visual search. *J Neurosci, 28*(24), 6141-6151. doi:10.1523/JNEUROSCI.1262-08.2008

Fagioli, S., & Macaluso, E. (2009). Attending to multiple visual streams: interactions between location-based and category-based attentional selection. *J Cogn Neurosci, 21*(8), 1628-1641. doi:10.1162/jocn.2009.21116

Fagioli, S., & Macaluso, E. (2016). Neural Correlates of Divided Attention in Natural Scenes. *J Cogn Neurosci, 28*(9), 1392-1405. doi:10.1162/jocn_a_00980

Fairhall, S. L., Indovina, I., Driver, J., & Macaluso, E. (2009). The brain network underlying serial visual search: comparing overt and covert spatial orienting, for activations and for effective connectivity. *Cereb Cortex, 19*(12), 2946-2958. doi:10.1093/cercor/bhp064

Fusser, F., Linden, D. E., Rahm, B., Hampel, H., Haenschel, C., & Mayer, J. S. (2011). Common capacity-limited neural mechanisms of selective attention and spatial working memory encoding. *Eur J Neurosci, 34*(5), 827-838. doi:10.1111/j.1460-9568.2011.07794.x

Galati, G., Committeri, G., Sanes, J. N., & Pizzamiglio, L. (2001). Spatial coding of visual and somatic sensory information in body-centred coordinates. *Eur J Neurosci, 14*(4), 737-746. doi:10.1046/j.0953-816x.2001.01674.x

Galati, G., Lobel, E., Vallar, G., Berthoz, A., Pizzamiglio, L., & Le Bihan, D. (2000). The neural basis of egocentric and allocentric coding of space in humans: a functional magnetic resonance study. *Exp Brain Res, 133*(2), 156-164. doi:10.1007/s002210000375

Geier, C. F., Garver, K. E., & Luna, B. (2007). Circuitry underlying temporally extended spatial working memory. *Neuroimage, 35*(2), 904-915. doi:10.1016/j.neuroimage.2006.12.022

Giesbrecht, B., Woldorff, M. G., Song, A. W., & Mangun, G. R. (2003). Neural mechanisms of top-down control during spatial and feature attention. *Neuroimage, 19*(3), 496-512. doi:10.1016/s1053-8119(03)00162-9

Gitelman, D. R., Nobre, A. C., Parrish, T. B., LaBar, K. S., Kim, Y. H., Meyer, J. R., & Mesulam, M. (1999). A large-scale distributed network for covert spatial attention: further anatomical delineation based on stringent behavioural and cognitive controls. *Brain, 122 ( Pt 6)*, 1093-1106. doi:10.1093/brain/122.6.1093

Halari, R., Sharma, T., Hines, M., Andrew, C., Simmons, A., & Kumari, V. (2006). Comparable fMRI activity with differential behavioural performance on mental rotation and overt verbal fluency tasks in healthy men and women. *Exp Brain Res, 169*(1), 1-14. doi:10.1007/s00221-005-0118-7

Harris, I. M., Egan, G. F., Sonkkila, C., Tochon-Danguy, H. J., Paxinos, G., & Watson, J. D. (2000). Selective right parietal lobe activation during mental rotation: a parametric PET study. *Brain, 123 ( Pt 1)*, 65-73. doi:10.1093/brain/123.1.65

Hautzel, H., Mottaghy, F. M., Schmidt, D., Zemb, M., Shah, N. J., Muller-Gartner, H. W., & Krause, B. J. (2002). Topographic segregation and convergence of verbal, object, shape and spatial working memory in humans. *Neurosci Lett, 323*(2), 156-160. doi:10.1016/s0304-3940(02)00125-8

Humphreys, G. F., & Lambon Ralph, M. A. (2017). Mapping Domain-Selective and Counterpointed Domain-General Higher Cognitive Functions in the Lateral Parietal Cortex: Evidence from fMRI Comparisons of Difficulty-Varying Semantic Versus Visuo-Spatial Tasks, and Functional Connectivity Analyses. *Cereb Cortex, 27*(8), 4199-4212. doi:10.1093/cercor/bhx107

Indovina, I., & Macaluso, E. (2007). Dissociation of stimulus relevance and saliency factors during shifts of visuospatial attention. *Cereb Cortex, 17*(7), 1701-1711. doi:10.1093/cercor/bhl081

Johnston, S., Leek, E. C., Atherton, C., Thacker, N., & Jackson, A. (2004). Functional contribution of medial premotor cortex to visuo-spatial transformation in humans. *Neurosci Lett, 355*(3), 209-212. doi:10.1016/j.neulet.2003.11.011

Jonides, J., Smith, E. E., Koeppe, R. A., Awh, E., Minoshima, S., & Mintun, M. A. (1993). Spatial working memory in humans as revealed by PET. *Nature, 363*(6430), 623-625. doi:10.1038/363623a0

Jordan, K., Heinze, H. J., Lutz, K., Kanowski, M., & Jancke, L. (2001). Cortical activations during the mental rotation of different visual objects. *Neuroimage, 13*(1), 143-152. doi:10.1006/nimg.2000.0677

Jordan, K., Wustenberg, T., Heinze, H. J., Peters, M., & Jancke, L. (2002). Women and men exhibit different cortical activation patterns during mental rotation tasks. *Neuropsychologia, 40*(13), 2397-2408. doi:10.1016/s0028-3932(02)00076-3

Kelley, T. A., Serences, J. T., Giesbrecht, B., & Yantis, S. (2008). Cortical mechanisms for shifting and holding visuospatial attention. *Cereb Cortex, 18*(1), 114-125. doi:10.1093/cercor/bhm036

Kim, Y. H., Gitelman, D. R., Nobre, A. C., Parrish, T. B., LaBar, K. S., & Mesulam, M. M. (1999). The large-scale neural network for spatial attention displays multifunctional overlap but differential asymmetry. *Neuroimage, 9*(3), 269-277. doi:10.1006/nimg.1999.0408

Kincade, J. M., Abrams, R. A., Astafiev, S. V., Shulman, G. L., & Corbetta, M. (2005). An event-related functional magnetic resonance imaging study of voluntary and stimulus-driven orienting of attention. *J Neurosci, 25*(18), 4593-4604. doi:10.1523/JNEUROSCI.0236-05.2005

Kosslyn, S. M., DiGirolamo, G. J., Thompson, W. L., & Alpert, N. M. (1998). Mental rotation of objects versus hands: neural mechanisms revealed by positron emission tomography. *Psychophysiology, 35*(2), 151-161.

Kosslyn, S. M., Thompson, W. L., Gitelman, D. R., & Alpert, N. M. (1998). Neural systems that encode categorical versus coordinate spatial relations: PET investigations. *Psychobiology, 26*(4), 333-347.

Krumbholz, K., Nobis, E. A., Weatheritt, R. J., & Fink, G. R. (2009). Executive control of spatial attention shifts in the auditory compared to the visual modality. *Hum Brain Mapp, 30*(5), 1457-1469. doi:10.1002/hbm.20615

LaBar, K. S., Gitelman, D. R., Parrish, T. B., & Mesulam, M. (1999). Neuroanatomic overlap of working memory and spatial attention networks: a functional MRI comparison within subjects. *Neuroimage, 10*(6), 695-704. doi:10.1006/nimg.1999.0503

Lamm, C., Windischberger, C., Leodolter, U., Moser, E., & Bauer, H. (2001). Evidence for premotor cortex activity during dynamic visuospatial imagery from single-trial functional magnetic resonance imaging and event-related slow cortical potentials. *Neuroimage, 14*(2), 268-283. doi:10.1006/nimg.2001.0850

Lamm, C., Windischberger, C., Moser, E., & Bauer, H. (2007). The functional role of dorso-lateral premotor cortex during mental rotation: an event-related fMRI study separating cognitive processing steps using a novel task paradigm. *Neuroimage, 36*(4), 1374-1386. doi:10.1016/j.neuroimage.2007.04.012

Lamp, G., Alexander, B., Laycock, R., Crewther, D. P., & Crewther, S. G. (2016). Mapping of the Underlying Neural Mechanisms of Maintenance and Manipulation in Visuo-Spatial Working Memory Using An n-back Mental Rotation Task: A Functional Magnetic Resonance Imaging Study. *Front Behav Neurosci, 10*, 87. doi:10.3389/fnbeh.2016.00087

Leek, E. C., Yuen, K. S., & Johnston, S. J. (2016). Domain General Sequence Operations Contribute to Pre-SMA Involvement in Visuo-spatial Processing. *Front Hum Neurosci, 10*, 9. doi:10.3389/fnhum.2016.00009

Lepsien, J., Griffin, I. C., Devlin, J. T., & Nobre, A. C. (2005). Directing spatial attention in mental representations: Interactions between attentional orienting and working-memory load. *Neuroimage, 26*(3), 733-743. doi:10.1016/j.neuroimage.2005.02.026

Leung, H. C., Seelig, D., & Gore, J. C. (2004). The effect of memory load on cortical activity in the spatial working memory circuit. *Cogn Affect Behav Neurosci, 4*(4), 553-563. doi:10.3758/cabn.4.4.553

Lipp, I., Benedek, M., Fink, A., Koschutnig, K., Reishofer, G., Bergner, S., . . . Neubauer, A. (2012). Investigating neural efficiency in the visuo-spatial domain: an FMRI study. *PLoS One, 7*(12), e51316. doi:10.1371/journal.pone.0051316

Macaluso, E., Eimer, M., Frith, C. D., & Driver, J. (2003). Preparatory states in crossmodal spatial attention: spatial specificity and possible control mechanisms. *Exp Brain Res, 149*(1), 62-74. doi:10.1007/s00221-002-1335-y

Macaluso, E., Frith, C. D., & Driver, J. (2002). Supramodal effects of covert spatial orienting triggered by visual or tactile events. *J Cogn Neurosci, 14*(3), 389-401. doi:10.1162/089892902317361912

Macaluso, E., & Patria, F. (2007). Spatial re-orienting of visual attention along the horizontal or the vertical axis. *Exp Brain Res, 180*(1), 23-34. doi:10.1007/s00221-006-0841-8

Manoach, D. S., White, N. S., Lindgren, K. A., Heckers, S., Coleman, M. J., Dubal, S., & Holzman, P. S. (2004). Hemispheric specialization of the lateral prefrontal cortex for strategic processing during spatial and shape working memory. *Neuroimage, 21*(3), 894-903. doi:10.1016/j.neuroimage.2003.10.025

Milivojevic, B., Hamm, J. P., & Corballis, M. C. (2009). Functional neuroanatomy of mental rotation. *J Cogn Neurosci, 21*(5), 945-959. doi:10.1162/jocn.2009.21085

Nardo, D., Santangelo, V., & Macaluso, E. (2011). Stimulus-driven orienting of visuo-spatial attention in complex dynamic environments. *Neuron, 69*(5), 1015-1028. doi:10.1016/j.neuron.2011.02.020

Natale, E., Marzi, C. A., & Macaluso, E. (2009). FMRI correlates of visuo-spatial reorienting investigated with an attention shifting double-cue paradigm. *Hum Brain Mapp, 30*(8), 2367-2381. doi:10.1002/hbm.20675

Nemmi, F., Boccia, M., Piccardi, L., Galati, G., & Guariglia, C. (2013). Segregation of neural circuits involved in spatial learning in reaching and navigational space. *Neuropsychologia, 51*(8), 1561-1570. doi:10.1016/j.neuropsychologia.2013.03.031

Ng, V. W., Bullmore, E. T., de Zubicaray, G. I., Cooper, A., Suckling, J., & Williams, S. C. (2001). Identifying rate-limiting nodes in large-scale cortical networks for visuospatial processing: an illustration using fMRI. *J Cogn Neurosci, 13*(4), 537-545. doi:10.1162/08989290152001943

Nobre, A. C., Sebestyen, G. N., Gitelman, D. R., Mesulam, M. M., Frackowiak, R. S., & Frith, C. D. (1997). Functional localization of the system for visuospatial attention using positron emission tomography. *Brain, 120 ( Pt 3)*, 515-533. doi:10.1093/brain/120.3.515

Nystrom, L. E., Braver, T. S., Sabb, F. W., Delgado, M. R., Noll, D. C., & Cohen, J. D. (2000). Working memory for letters, shapes, and locations: fMRI evidence against stimulus-based regional organization in human prefrontal cortex. *Neuroimage, 11*(5 Pt 1), 424-446. doi:10.1006/nimg.2000.0572

Ogawa, A., & Macaluso, E. (2015). Orienting of visuo-spatial attention in complex 3D space: Search and detection. *Hum Brain Mapp, 36*(6), 2231-2247. doi:10.1002/hbm.22767

Owen, A. M., Evans, A. C., & Petrides, M. (1996). Evidence for a two-stage model of spatial working memory processing within the lateral frontal cortex: a positron emission tomography study. *Cereb Cortex, 6*(1), 31-38. doi:10.1093/cercor/6.1.31

Peelen, M. V., Heslenfeld, D. J., & Theeuwes, J. (2004). Endogenous and exogenous attention shifts are mediated by the same large-scale neural network. *Neuroimage, 22*(2), 822-830. doi:10.1016/j.neuroimage.2004.01.044

Podzebenko, K., Egan, G. F., & Watson, J. D. (2002). Widespread dorsal stream activation during a parametric mental rotation task, revealed with functional magnetic resonance imaging. *Neuroimage, 15*(3), 547-558. doi:10.1006/nimg.2001.0999

Podzebenko, K., Egan, G. F., & Watson, J. D. (2005). Real and imaginary rotary motion processing: functional parcellation of the human parietal lobe revealed by fMRI. *J Cogn Neurosci, 17*(1), 24-36. doi:10.1162/0898929052879996

Ricciardi, E., Bonino, D., Gentili, C., Sani, L., Pietrini, P., & Vecchi, T. (2006). Neural correlates of spatial working memory in humans: a functional magnetic resonance imaging study comparing visual and tactile processes. *Neuroscience, 139*(1), 339-349. doi:10.1016/j.neuroscience.2005.08.045

Rosen, A. C., Rao, S. M., Caffarra, P., Scaglioni, A., Bobholz, J. A., Woodley, S. J., . . . Binder, J. R. (1999). Neural basis of endogenous and exogenous spatial orienting. A functional MRI study. *J Cogn Neurosci, 11*(2), 135-152. doi:10.1162/089892999563283

Rowe, J. B., Toni, I., Josephs, O., Frackowiak, R. S., & Passingham, R. E. (2000). The prefrontal cortex: response selection or maintenance within working memory? *Science, 288*(5471), 1656-1660. doi:10.1126/science.288.5471.1656

Sala, J. B., Rama, P., & Courtney, S. M. (2003). Functional topography of a distributed neural system for spatial and nonspatial information maintenance in working memory. *Neuropsychologia, 41*(3), 341-356. doi:10.1016/s0028-3932(02)00166-5

Salmi, J., Rinne, T., Degerman, A., Salonen, O., & Alho, K. (2007). Orienting and maintenance of spatial attention in audition and vision: multimodal and modality-specific brain activations. *Brain Struct Funct, 212*(2), 181-194. doi:10.1007/s00429-007-0152-2

Santangelo, V., Olivetti Belardinelli, M., Spence, C., & Macaluso, E. (2009). Interactions between voluntary and stimulus-driven spatial attention mechanisms across sensory modalities. *J Cogn Neurosci, 21*(12), 2384-2397. doi:10.1162/jocn.2008.21178

Schmidt, D., Krause, B. J., Weiss, P. H., Fink, G. R., Shah, N. J., Amorim, M. A., . . . Berthoz, A. (2007). Visuospatial working memory and changes of the point of view in 3D space. *Neuroimage, 36*(3), 955-968. doi:10.1016/j.neuroimage.2007.03.050

Schubotz, R. I., & von Cramon, D. Y. (2001). Functional organization of the lateral premotor cortex: fMRI reveals different regions activated by anticipation of object properties, location and speed. *Brain Res Cogn Brain Res, 11*(1), 97-112. doi:10.1016/s0926-6410(00)00069-0

Seurinck, R., Vingerhoets, G., de Lange, F. P., & Achten, E. (2004). Does egocentric mental rotation elicit sex differences? *Neuroimage, 23*(4), 1440-1449. doi:10.1016/j.neuroimage.2004.08.010

Seurinck, R., Vingerhoets, G., Vandemaele, P., Deblaere, K., & Achten, E. (2005). Trial pacing in mental rotation tasks. *Neuroimage, 25*(4), 1187-1196. doi:10.1016/j.neuroimage.2005.01.010

Seydell-Greenwald, A., Ferrara, K., Chambers, C. E., Newport, E. L., & Landau, B. (2017). Bilateral parietal activations for complex visual-spatial functions: Evidence from a visual-spatial construction task. *Neuropsychologia, 106*, 194-206. doi:10.1016/j.neuropsychologia.2017.10.005

Shen, L., Hu, X., Yacoub, E., & Ugurbil, K. (1999). Neural correlates of visual form and visual spatial processing. *Hum Brain Mapp, 8*(1), 60-71.

Simon, S. R., Meunier, M., Piettre, L., Berardi, A. M., Segebarth, C. M., & Boussaoud, D. (2002). Spatial attention and memory versus motor preparation: premotor cortex involvement as revealed by fMRI. *J Neurophysiol, 88*(4), 2047-2057. doi:10.1152/jn.2002.88.4.2047

Smith, D. V., Davis, B., Niu, K., Healy, E. W., Bonilha, L., Fridriksson, J., . . . Rorden, C. (2010). Spatial attention evokes similar activation patterns for visual and auditory stimuli. *J Cogn Neurosci, 22*(2), 347-361. doi:10.1162/jocn.2009.21241

Smith, E. E., Jonides, J., & Koeppe, R. A. (1996). Dissociating verbal and spatial working memory using PET. *Cereb Cortex, 6*(1), 11-20. doi:10.1093/cercor/6.1.11

Smith, E. E., Jonides, J., Koeppe, R. A., Awh, E., Schumacher, E. H., & Minoshima, S. (1995). Spatial versus Object Working Memory: PET Investigations. *J Cogn Neurosci, 7*(3), 337-356. doi:10.1162/jocn.1995.7.3.337

Summerfield, J. J., Lepsien, J., Gitelman, D. R., Mesulam, M. M., & Nobre, A. C. (2006). Orienting attention based on long-term memory experience. *Neuron, 49*(6), 905-916. doi:10.1016/j.neuron.2006.01.021

Tagaris, G. A., Kim, S. G., Strupp, J. P., Andersen, P., Ugurbil, K., & Georgopoulos, A. P. (1997). Mental rotation studied by functional magnetic resonance imaging at high field (4 tesla): performance and cortical activation. *J Cogn Neurosci, 9*(4), 419-432. doi:10.1162/jocn.1997.9.4.419

Tagaris, G. A., Richter, W., Kim, S. G., Pellizzer, G., Andersen, P., Ugurbil, K., & Georgopoulos, A. P. (1998). Functional magnetic resonance imaging of mental rotation and memory scanning: a multidimensional scaling analysis of brain activation patterns. *Brain Res Brain Res Rev, 26*(2-3), 106-112. doi:10.1016/s0165-0173(97)00060-x

Thiel, C. M., Zilles, K., & Fink, G. R. (2004). Cerebral correlates of alerting, orienting and reorienting of visuospatial attention: an event-related fMRI study. *Neuroimage, 21*(1), 318-328. doi:10.1016/j.neuroimage.2003.08.044

Thomason, M. E., Race, E., Burrows, B., Whitfield-Gabrieli, S., Glover, G. H., & Gabrieli, J. D. (2009). Development of spatial and verbal working memory capacity in the human brain. *J Cogn Neurosci, 21*(2), 316-332. doi:10.1162/jocn.2008.21028

Thomsen, T., Hugdahl, K., Ersland, L., Barndon, R., Lundervold, A., Smievoll, A. I., . . . Sundberg, H. (2000). Functional magnetic resonance imaging (fMRI) study of sex differences in a mental rotation task. *Med Sci Monit, 6*(6), 1186-1196.

Umla-Runge, K., Zimmer, H. D., Krick, C. M., & Reith, W. (2011). fMRI correlates of working memory: specific posterior representation sites for motion and position information. *Brain Res, 1382*, 206-218. doi:10.1016/j.brainres.2011.01.052

Vallar, G., Lobel, E., Galati, G., Berthoz, A., Pizzamiglio, L., & Le Bihan, D. (1999). A fronto-parietal system for computing the egocentric spatial frame of reference in humans. *Exp Brain Res, 124*(3), 281-286. doi:10.1007/s002210050624

Vandenberghe, R., Gitelman, D. R., Parrish, T. B., & Mesulam, M. M. (2001). Functional specificity of superior parietal mediation of spatial shifting. *Neuroimage, 14*(3), 661-673. doi:10.1006/nimg.2001.0860

Vanrie, J., Beatse, E., Wagemans, J., Sunaert, S., & Van Hecke, P. (2002). Mental rotation versus invariant features in object perception from different viewpoints: an fMRI study. *Neuropsychologia, 40*(7), 917-930. doi:10.1016/s0028-3932(01)00161-0

Ventre-Dominey, J., Bailly, A., Lavenne, F., Lebars, D., Mollion, H., Costes, N., & Dominey, P. F. (2005). Double dissociation in neural correlates of visual working memory: a PET study. *Brain Res Cogn Brain Res, 25*(3), 747-759. doi:10.1016/j.cogbrainres.2005.09.004

Vingerhoets, G., de Lange, F. P., Vandemaele, P., Deblaere, K., & Achten, E. (2002). Motor imagery in mental rotation: an fMRI study. *Neuroimage, 17*(3), 1623-1633. doi:10.1006/nimg.2002.1290

Walter, H., Bretschneider, V., Gron, G., Zurowski, B., Wunderlich, A. P., Tomczak, R., & Spitzer, M. (2003). Evidence for quantitative domain dominance for verbal and spatial working memory in frontal and parietal cortex. *Cortex, 39*(4-5), 897-911. doi:10.1016/s0010-9452(08)70869-4

Weeks, R. A., Aziz-Sultan, A., Bushara, K. O., Tian, B., Wessinger, C. M., Dang, N., . . . Hallett, M. (1999). A PET study of human auditory spatial processing. *Neurosci Lett, 262*(3), 155-158. doi:10.1016/s0304-3940(99)00062-2

Weiss, E., Siedentopf, C. M., Hofer, A., Deisenhammer, E. A., Hoptman, M. J., Kremser, C., . . . Delazer, M. (2003). Sex differences in brain activation pattern during a visuospatial cognitive task: a functional magnetic resonance imaging study in healthy volunteers. *Neurosci Lett, 344*(3), 169-172. doi:10.1016/s0304-3940(03)00406-3

Windischberger, C., Lamm, C., Bauer, H., & Moser, E. (2003). Human motor cortex activity during mental rotation. *Neuroimage, 20*(1), 225-232. doi:10.1016/s1053-8119(03)00235-0

Wraga, M., Thompson, W. L., Alpert, N. M., & Kosslyn, S. M. (2003). Implicit transfer of motor strategies in mental rotation. *Brain Cogn, 52*(2), 135-143. doi:10.1016/s0278-2626(03)00033-2

Yantis, S., Schwarzbach, J., Serences, J. T., Carlson, R. L., Steinmetz, M. A., Pekar, J. J., & Courtney, S. M. (2002). Transient neural activity in human parietal cortex during spatial attention shifts. *Nat Neurosci, 5*(10), 995-1002. doi:10.1038/nn921

Zacks, J., Rypma, B., Gabrieli, J. D., Tversky, B., & Glover, G. H. (1999). Imagined transformations of bodies: an fMRI investigation. *Neuropsychologia, 37*(9), 1029-1040. doi:10.1016/s0028-3932(99)00012-3

Zaehle, T., Jordan, K., Wustenberg, T., Baudewig, J., Dechent, P., & Mast, F. W. (2007). The neural basis of the egocentric and allocentric spatial frame of reference. *Brain Res, 1137*(1), 92-103. doi:10.1016/j.brainres.2006.12.044
