## Supplementary file D for "From *ATOM* to *GradiATOM*: Cortical gradients support time and space processing as revealed by a meta-analysis of neuroimaging studies"

**Supplementary Information D**

**List of Included Studies for the “Time” meta-analysis**

(Apaydin et al., 2018; Aso, Hanakawa, Aso, & Fukuyama, 2010; Basso, Nichelli, Wharton, Peterson, & Grafman, 2003; Belin et al., 2002; Bengtsson, Ehrsson, Forssberg, & Ullen, 2004, 2005; Bengtsson et al., 2009; Beudel, Renken, Leenders, & de Jong, 2009; Bijsterbosch et al., 2011; Bortoletto & Cunnington, 2010; Brendel et al., 2010; Brunia, de Jong, van den Berg-Lenssen, & Paans, 2000; Bueti & Macaluso, 2010, 2011; Bueti, Walsh, Frith, & Rees, 2008; Carvalho, Chaim, Sanchez, & de Araujo, 2016; Cerasa, Hagberg, Bianciardi, & Sabatini, 2005; Chen, Penhune, & Zatorre, 2008; Chen, Zatorre, & Penhune, 2006; Coull, Charras, Donadieu, Droit-Volet, & Vidal, 2015; Coull, Davranche, Nazarian, & Vidal, 2013; Coull, Hwang, Leyton, & Dagher, 2012; Coull, Nazarian, & Vidal, 2008; Coull & Nobre, 1998; Coull, Vidal, Goulon, Nazarian, & Craig, 2008; Coull, Vidal, Nazarian, & Macar, 2004; Cunnington, Windischberger, Deecke, & Moser, 2002; Davranche, Nazarian, Vidal, & Coull, 2011; Ferrandez et al., 2003; Filip, Losak, Kasparek, Vanicek, & Bares, 2016; Gandour et al., 2002; Grahn & Brett, 2007; Grahn, Henry, & McAuley, 2011; Grahn & McAuley, 2009; Grahn & Rowe, 2009, 2013; Gutyrchik et al., 2010; Harrington et al., 2004; Harrington, Zimbelman, Hinton, & Rao, 2010; Hayashi et al., 2015; Henry, Herrmann, & Obleser, 2015; Jahanshahi, Jones, Dirnberger, & Frith, 2006; Jancke, Loose, Lutz, Specht, & Shah, 2000; Jantzen, Oullier, Marshall, Steinberg, & Kelso, 2007; Jantzen, Steinberg, & Kelso, 2004, 2005; Jech, Dusek, Wackermann, & Vymazal, 2005; Jeuptner, Flerich, Weiller, Mueller, & Diener, 1996; Karabanov, Blom, Forsman, & Ullen, 2009; Kawashima et al., 1999; Kawashima et al., 2000; Konoike et al., 2015; Konoike et al., 2012; Kudo et al., 2004; Larsson, Gulyas, & Roland, 1996; Lejeune et al., 1997; Lewis & Miall, 2002, 2003; Lewis, Wing, Pope, Praamstra, & Miall, 2004; C. Li, Chen, Han, Chui, & Wu, 2012; Y. Li, Mo, & Chen, 2015; Livesey, Wall, & Smith, 2007; Lutz, Specht, Shah, & Jancke, 2000; Macar, Anton, Bonnet, & Vidal, 2004; Macar et al., 2002; Maquet et al., 1996; Marchant & Driver, 2013; Mayville, Jantzen, Fuchs, Steinberg, & Kelso, 2002; Morillon, Kell, & Giraud, 2009; Nenadic et al., 2003; Neufang, Fink, Herpertz-Dahlmann, Willmes, & Konrad, 2008; O'Reilly, Mesulam, & Nobre, 2008; Ortuno et al., 2002; Oullier, Jantzen, Steinberg, & Kelso, 2005; Pastor, Macaluso, Day, & Frackowiak, 2006; Penhune & Doyon, 2002; Penhune, Zattore, & Evans, 1998; Pfeuty, Dilharreguy, Gerlier, & Allard, 2015; Pope, Wing, Praamstra, & Miall, 2005; Pouthas et al., 2005; Ramnani & Passingham, 2001; Rao et al., 1997; Rao, Mayer, & Harrington, 2001; Riecker, Wildgruber, Mathiak, Grodd, & Ackermann, 2003; Sakai et al., 1999; Sakai et al., 2000; Sakai, Ramnani, & Passingham, 2002; Schubotz, Friederici, & von Cramon, 2000; Schubotz & von Cramon, 2001; Schubotz, von Cramon, & Lohmann, 2003; Shergill, Tracy, Seal, Rubia, & McGuire, 2006; Shih, Chen, et al., 2009; Shih, Kuo, Yeh, Tzeng, & Hsieh, 2009; Skagerlund, Karlsson, & Traff, 2016; A. Smith, Taylor, Lidzba, & Rubia, 2003; A. B. Smith et al., 2011; Taniwaki et al., 2006; Teki, Grube, Kumar, & Griffiths, 2011; Thaut, Demartin, & Sanes, 2008; Tipples, Brattan, & Johnston, 2013; Tomasi, Wang, Studentsova, & Volkow, 2015; Tracy, Faro, Mohamed, Pinsk, & Pinus, 2000; Tregellas, Davalos, & Rojas, 2006; Ustun, Kale, & Cicek, 2017; Wiener, Lee, Lohoff, & Coslett, 2014; Wittmann, Simmons, Aron, & Paulus, 2010; Wittmann et al., 2011; Wittmann, van Wassenhove, Craig, & Paulus, 2010; Woods et al., 2014)

Apaydin, N., Ustun, S., Kale, E. H., Celikag, I., Ozguven, H. D., Baskak, B., & Cicek, M. (2018). Neural Mechanisms Underlying Time Perception and Reward Anticipation. *Front Hum Neurosci, 12*, 115. doi:10.3389/fnhum.2018.00115

Aso, K., Hanakawa, T., Aso, T., & Fukuyama, H. (2010). Cerebro-cerebellar interactions underlying temporal information processing. *J Cogn Neurosci, 22*(12), 2913-2925. doi:10.1162/jocn.2010.21429

Basso, G., Nichelli, P., Wharton, C. M., Peterson, M., & Grafman, J. (2003). Distributed neural systems for temporal production: a functional MRI study. *Brain Res Bull, 59*(5), 405-411. doi:10.1016/s0361-9230(02)00941-3

Belin, P., McAdams, S., Thivard, L., Smith, B., Savel, S., Zilbovicius, M., . . . Samson, Y. (2002). The neuroanatomical substrate of sound duration discrimination. *Neuropsychologia, 40*(12), 1956-1964. doi:10.1016/s0028-3932(02)00062-3

Bengtsson, S. L., Ehrsson, H. H., Forssberg, H., & Ullen, F. (2004). Dissociating brain regions controlling the temporal and ordinal structure of learned movement sequences. *Eur J Neurosci, 19*(9), 2591-2602. doi:10.1111/j.0953-816X.2004.03269.x

Bengtsson, S. L., Ehrsson, H. H., Forssberg, H., & Ullen, F. (2005). Effector-independent voluntary timing: behavioural and neuroimaging evidence. *Eur J Neurosci, 22*(12), 3255-3265. doi:10.1111/j.1460-9568.2005.04517.x

Bengtsson, S. L., Ullen, F., Ehrsson, H. H., Hashimoto, T., Kito, T., Naito, E., . . . Sadato, N. (2009). Listening to rhythms activates motor and premotor cortices. *Cortex, 45*(1), 62-71. doi:10.1016/j.cortex.2008.07.002

Beudel, M., Renken, R., Leenders, K. L., & de Jong, B. M. (2009). Cerebral representations of space and time. *Neuroimage, 44*(3), 1032-1040. doi:10.1016/j.neuroimage.2008.09.028

Bijsterbosch, J. D., Lee, K. H., Hunter, M. D., Tsoi, D. T., Lankappa, S., Wilkinson, I. D., . . . Woodruff, P. W. (2011). The role of the cerebellum in sub- and supraliminal error correction during sensorimotor synchronization: evidence from fMRI and TMS. *J Cogn Neurosci, 23*(5), 1100-1112. doi:10.1162/jocn.2010.21506

Bortoletto, M., & Cunnington, R. (2010). Motor timing and motor sequencing contribute differently to the preparation for voluntary movement. *Neuroimage, 49*(4), 3338-3348. doi:10.1016/j.neuroimage.2009.11.048

Brendel, B., Hertrich, I., Erb, M., Lindner, A., Riecker, A., Grodd, W., & Ackermann, H. (2010). The contribution of mesiofrontal cortex to the preparation and execution of repetitive syllable productions: an fMRI study. *Neuroimage, 50*(3), 1219-1230. doi:10.1016/j.neuroimage.2010.01.039

Brunia, C. H., de Jong, B. M., van den Berg-Lenssen, M. M., & Paans, A. M. (2000). Visual feedback about time estimation is related to a right hemisphere activation measured by PET. *Exp Brain Res, 130*(3), 328-337. doi:10.1007/s002219900293

Bueti, D., & Macaluso, E. (2010). Auditory temporal expectations modulate activity in visual cortex. *Neuroimage, 51*(3), 1168-1183. doi:10.1016/j.neuroimage.2010.03.023

Bueti, D., & Macaluso, E. (2011). Physiological correlates of subjective time: evidence for the temporal accumulator hypothesis. *Neuroimage, 57*(3), 1251-1263. doi:10.1016/j.neuroimage.2011.05.014

Bueti, D., Walsh, V., Frith, C., & Rees, G. (2008). Different brain circuits underlie motor and perceptual representations of temporal intervals. *J Cogn Neurosci, 20*(2), 204-214. doi:10.1162/jocn.2008.20017

Carvalho, F. M., Chaim, K. T., Sanchez, T. A., & de Araujo, D. B. (2016). Time-Perception Network and Default Mode Network Are Associated with Temporal Prediction in a Periodic Motion Task. *Front Hum Neurosci, 10*, 268. doi:10.3389/fnhum.2016.00268

Cerasa, A., Hagberg, G. E., Bianciardi, M., & Sabatini, U. (2005). Visually cued motor synchronization: modulation of fMRI activation patterns by baseline condition. *Neurosci Lett, 373*(1), 32-37. doi:10.1016/j.neulet.2004.09.076

Chen, J. L., Penhune, V. B., & Zatorre, R. J. (2008). Listening to musical rhythms recruits motor regions of the brain. *Cereb Cortex, 18*(12), 2844-2854. doi:10.1093/cercor/bhn042

Chen, J. L., Zatorre, R. J., & Penhune, V. B. (2006). Interactions between auditory and dorsal premotor cortex during synchronization to musical rhythms. *Neuroimage, 32*(4), 1771-1781. doi:10.1016/j.neuroimage.2006.04.207

Coull, J. T., Charras, P., Donadieu, M., Droit-Volet, S., & Vidal, F. (2015). SMA Selectively Codes the Active Accumulation of Temporal, Not Spatial, Magnitude. *J Cogn Neurosci, 27*(11), 2281-2298. doi:10.1162/jocn_a_00854

Coull, J. T., Davranche, K., Nazarian, B., & Vidal, F. (2013). Functional anatomy of timing differs for production versus prediction of time intervals. *Neuropsychologia, 51*(2), 309-319. doi:10.1016/j.neuropsychologia.2012.08.017

Coull, J. T., Hwang, H. J., Leyton, M., & Dagher, A. (2012). Dopamine precursor depletion impairs timing in healthy volunteers by attenuating activity in putamen and supplementary motor area. *J Neurosci, 32*(47), 16704-16715. doi:10.1523/JNEUROSCI.1258-12.2012

Coull, J. T., Nazarian, B., & Vidal, F. (2008). Timing, storage, and comparison of stimulus duration engage discrete anatomical components of a perceptual timing network. *J Cogn Neurosci, 20*(12), 2185-2197. doi:10.1162/jocn.2008.20153

Coull, J. T., & Nobre, A. C. (1998). Where and when to pay attention: the neural systems for directing attention to spatial locations and to time intervals as revealed by both PET and fMRI. *J Neurosci, 18*(18), 7426-7435.

Coull, J. T., Vidal, F., Goulon, C., Nazarian, B., & Craig, C. (2008). Using time-to-contact information to assess potential collision modulates both visual and temporal prediction networks. *Front Hum Neurosci, 2*, 10. doi:10.3389/neuro.09.010.2008

Coull, J. T., Vidal, F., Nazarian, B., & Macar, F. (2004). Functional anatomy of the attentional modulation of time estimation. *Science, 303*(5663), 1506-1508. doi:10.1126/science.1091573

Cunnington, R., Windischberger, C., Deecke, L., & Moser, E. (2002). The preparation and execution of self-initiated and externally-triggered movement: a study of event-related fMRI. *Neuroimage, 15*(2), 373-385. doi:10.1006/nimg.2001.0976

Davranche, K., Nazarian, B., Vidal, F., & Coull, J. (2011). Orienting attention in time activates left intraparietal sulcus for both perceptual and motor task goals. *J Cogn Neurosci, 23*(11), 3318-3330. doi:10.1162/jocn_a_00030

Ferrandez, A. M., Hugueville, L., Lehericy, S., Poline, J. B., Marsault, C., & Pouthas, V. (2003). Basal ganglia and supplementary motor area subtend duration perception: an fMRI study. *Neuroimage, 19*(4), 1532-1544. doi:10.1016/s1053-8119(03)00159-9

Filip, P., Losak, J., Kasparek, T., Vanicek, J., & Bares, M. (2016). Neural Network of Predictive Motor Timing in the Context of Gender Differences. *Neural Plast, 2016*, 2073454. doi:10.1155/2016/2073454

Gandour, J., Wong, D., Lowe, M., Dzemidzic, M., Satthamnuwong, N., Tong, Y., & Lurito, J. (2002). Neural circuitry underlying perception of duration depends on language experience. *Brain Lang, 83*(2), 268-290. doi:10.1016/s0093-934x(02)00033-0

Grahn, J. A., & Brett, M. (2007). Rhythm and beat perception in motor areas of the brain. *J Cogn Neurosci, 19*(5), 893-906. doi:10.1162/jocn.2007.19.5.893

Grahn, J. A., Henry, M. J., & McAuley, J. D. (2011). FMRI investigation of cross-modal interactions in beat perception: audition primes vision, but not vice versa. *Neuroimage, 54*(2), 1231-1243. doi:10.1016/j.neuroimage.2010.09.033

Grahn, J. A., & McAuley, J. D. (2009). Neural bases of individual differences in beat perception. *Neuroimage, 47*(4), 1894-1903. doi:10.1016/j.neuroimage.2009.04.039

Grahn, J. A., & Rowe, J. B. (2009). Feeling the beat: premotor and striatal interactions in musicians and nonmusicians during beat perception. *J Neurosci, 29*(23), 7540-7548. doi:10.1523/JNEUROSCI.2018-08.2009

Grahn, J. A., & Rowe, J. B. (2013). Finding and feeling the musical beat: striatal dissociations between detection and prediction of regularity. *Cereb Cortex, 23*(4), 913-921. doi:10.1093/cercor/bhs083

Gutyrchik, E., Churan, J., Meindl, T., Bokde, A. L., von Bernewitz, H., Born, C., . . . Wittmann, M. (2010). Functional neuroimaging of duration discrimination on two different time scales. *Neurosci Lett, 469*(3), 411-415. doi:10.1016/j.neulet.2009.12.040

Harrington, D. L., Boyd, L. A., Mayer, A. R., Sheltraw, D. M., Lee, R. R., Huang, M., & Rao, S. M. (2004). Neural representation of interval encoding and decision making. *Brain Res Cogn Brain Res, 21*(2), 193-205. doi:10.1016/j.cogbrainres.2004.01.010

Harrington, D. L., Zimbelman, J. L., Hinton, S. C., & Rao, S. M. (2010). Neural modulation of temporal encoding, maintenance, and decision processes. *Cereb Cortex, 20*(6), 1274-1285. doi:10.1093/cercor/bhp194

Hayashi, M. J., Ditye, T., Harada, T., Hashiguchi, M., Sadato, N., Carlson, S., . . . Kanai, R. (2015). Time Adaptation Shows Duration Selectivity in the Human Parietal Cortex. *PLoS Biol, 13*(9), e1002262. doi:10.1371/journal.pbio.1002262

Henry, M. J., Herrmann, B., & Obleser, J. (2015). Selective attention to temporal features on nested time scales. *Cereb Cortex, 25*(2), 450-459. doi:10.1093/cercor/bht240

Jahanshahi, M., Jones, C. R., Dirnberger, G., & Frith, C. D. (2006). The substantia nigra pars compacta and temporal processing. *J Neurosci, 26*(47), 12266-12273. doi:10.1523/JNEUROSCI.2540-06.2006

Jancke, L., Loose, R., Lutz, K., Specht, K., & Shah, N. J. (2000). Cortical activations during paced finger-tapping applying visual and auditory pacing stimuli. *Brain Res Cogn Brain Res, 10*(1-2), 51-66. doi:10.1016/s0926-6410(00)00022-7

Jantzen, K. J., Oullier, O., Marshall, M., Steinberg, F. L., & Kelso, J. A. (2007). A parametric fMRI investigation of context effects in sensorimotor timing and coordination. *Neuropsychologia, 45*(4), 673-684. doi:10.1016/j.neuropsychologia.2006.07.020

Jantzen, K. J., Steinberg, F. L., & Kelso, J. A. (2004). Brain networks underlying human timing behavior are influenced by prior context. *Proc Natl Acad Sci U S A, 101*(17), 6815-6820. doi:10.1073/pnas.0401300101

Jantzen, K. J., Steinberg, F. L., & Kelso, J. A. (2005). Functional MRI reveals the existence of modality and coordination-dependent timing networks. *Neuroimage, 25*(4), 1031-1042. doi:10.1016/j.neuroimage.2004.12.029

Jech, R., Dusek, P., Wackermann, J., & Vymazal, J. (2005). Cumulative blood oxygenation-level-dependent signal changes support the 'time accumulator' hypothesis. *Neuroreport, 16*(13), 1467-1471. doi:10.1097/01.wnr.0000175616.00936.1c

Jeuptner, M., Flerich, L., Weiller, C., Mueller, S. P., & Diener, H. C. (1996). The human cerebellum and temporal information processing—results from a PET experiment. *Neuroreport, 7*, 2761-2765.

Karabanov, A., Blom, O., Forsman, L., & Ullen, F. (2009). The dorsal auditory pathway is involved in performance of both visual and auditory rhythms. *Neuroimage, 44*(2), 480-488. doi:10.1016/j.neuroimage.2008.08.047

Kawashima, R., Inoue, K., Sugiura, M., Okada, K., Ogawa, A., & Fukuda, H. (1999). A positron emission tomography study of self-paced finger movements at different frequencies. *Neuroscience, 92*(1), 107-112. doi:10.1016/s0306-4522(98)00744-1

Kawashima, R., Okuda, J., Umetsu, A., Sugiura, M., Inoue, K., Suzuki, K., . . . Yamadori, A. (2000). Human cerebellum plays an important role in memory-timed finger movement: an fMRI study. *J Neurophysiol, 83*(2), 1079-1087. doi:10.1152/jn.2000.83.2.1079

Konoike, N., Kotozaki, Y., Jeong, H., Miyazaki, A., Sakaki, K., Shinada, T., . . . Nakamura, K. (2015). Temporal and Motor Representation of Rhythm in Fronto-Parietal Cortical Areas: An fMRI Study. *PLoS One, 10*(6), e0130120. doi:10.1371/journal.pone.0130120

Konoike, N., Kotozaki, Y., Miyachi, S., Miyauchi, C. M., Yomogida, Y., Akimoto, Y., . . . Nakamura, K. (2012). Rhythm information represented in the fronto-parieto-cerebellar motor system. *Neuroimage, 63*(1), 328-338. doi:10.1016/j.neuroimage.2012.07.002

Kudo, K., Miyazaki, M., Kimura, T., Yamanaka, K., Kadota, H., Hirashima, M., . . . Ohtsuki, T. (2004). Selective activation and deactivation of the human brain structures between speeded and precisely timed tapping responses to identical visual stimulus: an fMRI study. *Neuroimage, 22*(3), 1291-1301. doi:10.1016/j.neuroimage.2004.03.043

Larsson, J., Gulyas, B., & Roland, P. E. (1996). Cortical representation of self-paced finger movement. *Neuroreport, 7*(2), 463-468. doi:10.1097/00001756-199601310-00021

Lejeune, H., Maquet, P., Bonnet, M., Casini, L., Ferrara, A., Macar, F., . . . Vidal, F. (1997). The basic pattern of activation in motor and sensory temporal tasks: positron emission tomography data. *Neurosci Lett, 235*(1-2), 21-24. doi:10.1016/s0304-3940(97)00698-8

Lewis, P. A., & Miall, R. C. (2002). Brain activity during non-automatic motor production of discrete multi-second intervals. *Neuroreport, 13*(14), 1731-1735. doi:10.1097/00001756-200210070-00008

Lewis, P. A., & Miall, R. C. (2003). Brain activation patterns during measurement of sub- and supra-second intervals. *Neuropsychologia, 41*(12), 1583-1592. doi:10.1016/s0028-3932(03)00118-0

Lewis, P. A., Wing, A. M., Pope, P. A., Praamstra, P., & Miall, R. C. (2004). Brain activity correlates differentially with increasing temporal complexity of rhythms during initialisation, synchronisation, and continuation phases of paced finger tapping. *Neuropsychologia, 42*(10), 1301-1312. doi:10.1016/j.neuropsychologia.2004.03.001

Li, C., Chen, K., Han, H., Chui, D., & Wu, J. (2012). An FMRI study of the neural systems involved in visually cued auditory top-down spatial and temporal attention. *PLoS One, 7*(11), e49948. doi:10.1371/journal.pone.0049948

Li, Y., Mo, L., & Chen, Q. (2015). Differential contribution of velocity and distance to time estimation during self-initiated time-to-collision judgment. *Neuropsychologia, 73*, 35-47. doi:10.1016/j.neuropsychologia.2015.04.017

Livesey, A. C., Wall, M. B., & Smith, A. T. (2007). Time perception: manipulation of task difficulty dissociates clock functions from other cognitive demands. *Neuropsychologia, 45*(2), 321-331. doi:10.1016/j.neuropsychologia.2006.06.033

Lutz, K., Specht, K., Shah, N. J., & Jancke, L. (2000). Tapping movements according to regular and irregular visual timing signals investigated with fMRI. *Neuroreport, 11*(6), 1301-1306. doi:10.1097/00001756-200004270-00031

Macar, F., Anton, J. L., Bonnet, M., & Vidal, F. (2004). Timing functions of the supplementary motor area: an event-related fMRI study. *Brain Res Cogn Brain Res, 21*(2), 206-215. doi:10.1016/j.cogbrainres.2004.01.005

Macar, F., Lejeune, H., Bonnet, M., Ferrara, A., Pouthas, V., Vidal, F., & Maquet, P. (2002). Activation of the supplementary motor area and of attentional networks during temporal processing. *Exp Brain Res, 142*(4), 475-485. doi:10.1007/s00221-001-0953-0

Maquet, P., Lejeune, H., Pouthas, V., Bonnet, M., Casini, L., Macar, F., . . . Comar, D. (1996). Brain activation induced by estimation of duration: a PET study. *Neuroimage, 3*(2), 119-126. doi:10.1006/nimg.1996.0014

Marchant, J. L., & Driver, J. (2013). Visual and audiovisual effects of isochronous timing on visual perception and brain activity. *Cereb Cortex, 23*(6), 1290-1298. doi:10.1093/cercor/bhs095

Mayville, J. M., Jantzen, K. J., Fuchs, A., Steinberg, F. L., & Kelso, J. A. (2002). Cortical and subcortical networks underlying syncopated and synchronized coordination revealed using fMRI. Functional magnetic resonance imaging. *Hum Brain Mapp, 17*(4), 214-229. doi:10.1002/hbm.10065

Morillon, B., Kell, C. A., & Giraud, A. L. (2009). Three stages and four neural systems in time estimation. *J Neurosci, 29*(47), 14803-14811. doi:10.1523/JNEUROSCI.3222-09.2009

Nenadic, I., Gaser, C., Volz, H. P., Rammsayer, T., Hager, F., & Sauer, H. (2003). Processing of temporal information and the basal ganglia: new evidence from fMRI. *Exp Brain Res, 148*(2), 238-246. doi:10.1007/s00221-002-1188-4

Neufang, S., Fink, G. R., Herpertz-Dahlmann, B., Willmes, K., & Konrad, K. (2008). Developmental changes in neural activation and psychophysiological interaction patterns of brain regions associated with interference control and time perception. *Neuroimage, 43*(2), 399-409. doi:10.1016/j.neuroimage.2008.07.039

O'Reilly, J. X., Mesulam, M. M., & Nobre, A. C. (2008). The cerebellum predicts the timing of perceptual events. *J Neurosci, 28*(9), 2252-2260. doi:10.1523/JNEUROSCI.2742-07.2008

Ortuno, F., Ojeda, N., Arbizu, J., Lopez, P., Marti-Climent, J. M., Penuelas, I., & Cervera, S. (2002). Sustained attention in a counting task: normal performance and functional neuroanatomy. *Neuroimage, 17*(1), 411-420. doi:10.1006/nimg.2002.1168

Oullier, O., Jantzen, K. J., Steinberg, F. L., & Kelso, J. A. (2005). Neural substrates of real and imagined sensorimotor coordination. *Cereb Cortex, 15*(7), 975-985. doi:10.1093/cercor/bhh198

Pastor, M. A., Macaluso, E., Day, B. L., & Frackowiak, R. S. (2006). The neural basis of temporal auditory discrimination. *Neuroimage, 30*(2), 512-520. doi:10.1016/j.neuroimage.2005.09.053

Penhune, V. B., & Doyon, J. (2002). Dynamic cortical and subcortical networks in learning and delayed recall of timed motor sequences. *J Neurosci, 22*(4), 1397-1406.

Penhune, V. B., Zattore, R. J., & Evans, A. C. (1998). Cerebellar contributions to motor timing: a PET study of auditory and visual rhythm reproduction. *J Cogn Neurosci, 10*(6), 752-765. doi:10.1162/089892998563149

Pfeuty, M., Dilharreguy, B., Gerlier, L., & Allard, M. (2015). fMRI identifies the right inferior frontal cortex as the brain region where time interval processing is altered by negative emotional arousal. *Hum Brain Mapp, 36*(3), 981-995. doi:10.1002/hbm.22680

Pope, P., Wing, A. M., Praamstra, P., & Miall, R. C. (2005). Force related activations in rhythmic sequence production. *Neuroimage, 27*(4), 909-918. doi:10.1016/j.neuroimage.2005.05.010

Pouthas, V., George, N., Poline, J. B., Pfeuty, M., Vandemoorteele, P. F., Hugueville, L., . . . Renault, B. (2005). Neural network involved in time perception: an fMRI study comparing long and short interval estimation. *Hum Brain Mapp, 25*(4), 433-441. doi:10.1002/hbm.20126

Ramnani, N., & Passingham, R. E. (2001). Changes in the human brain during rhythm learning. *J Cogn Neurosci, 13*(7), 952-966. doi:10.1162/089892901753165863

Rao, S. M., Harrington, D. L., Haaland, K. Y., Bobholz, J. A., Cox, R. W., & Binder, J. R. (1997). Distributed neural systems underlying the timing of movements. *J Neurosci, 17*(14), 5528-5535.

Rao, S. M., Mayer, A. R., & Harrington, D. L. (2001). The evolution of brain activation during temporal processing. *Nat Neurosci, 4*(3), 317-323. doi:10.1038/85191

Riecker, A., Wildgruber, D., Mathiak, K., Grodd, W., & Ackermann, H. (2003). Parametric analysis of rate-dependent hemodynamic response functions of cortical and subcortical brain structures during auditorily cued finger tapping: a fMRI study. *Neuroimage, 18*(3), 731-739. doi:10.1016/s1053-8119(03)00003-x

Sakai, K., Hikosaka, O., Miyauchi, S., Takino, R., Tamada, T., Iwata, N. K., & Nielsen, M. (1999). Neural representation of a rhythm depends on its interval ratio. *J Neurosci, 19*(22), 10074-10081.

Sakai, K., Hikosaka, O., Takino, R., Miyauchi, S., Nielsen, M., & Tamada, T. (2000). What and when: parallel and convergent processing in motor control. *J Neurosci, 20*(7), 2691-2700.

Sakai, K., Ramnani, N., & Passingham, R. E. (2002). Learning of sequences of finger movements and timing: frontal lobe and action-oriented representation. *J Neurophysiol, 88*(4), 2035-2046. doi:10.1152/jn.2002.88.4.2035

Schubotz, R. I., Friederici, A. D., & von Cramon, D. Y. (2000). Time perception and motor timing: a common cortical and subcortical basis revealed by fMRI. *Neuroimage, 11*(1), 1-12. doi:10.1006/nimg.1999.0514

Schubotz, R. I., & von Cramon, D. Y. (2001). Interval and ordinal properties of sequences are associated with distinct premotor areas. *Cereb Cortex, 11*(3), 210-222. doi:10.1093/cercor/11.3.210

Schubotz, R. I., von Cramon, D. Y., & Lohmann, G. (2003). Auditory what, where, and when: a sensory somatotopy in lateral premotor cortex. *Neuroimage, 20*(1), 173-185. doi:10.1016/s1053-8119(03)00218-0

Shergill, S. S., Tracy, D. K., Seal, M., Rubia, K., & McGuire, P. (2006). Timing of covert articulation: an fMRI study. *Neuropsychologia, 44*(12), 2573-2577. doi:10.1016/j.neuropsychologia.2006.04.005

Shih, L. Y., Chen, L. F., Kuo, W. J., Yeh, T. C., Wu, Y. T., Tzeng, O. J., & Hsieh, J. C. (2009). Sensory acquisition in the cerebellum: an FMRI study of cerebrocerebellar interaction during visual duration discrimination. *Cerebellum, 8*(2), 116-126. doi:10.1007/s12311-008-0082-4

Shih, L. Y., Kuo, W. J., Yeh, T. C., Tzeng, O. J., & Hsieh, J. C. (2009). Common neural mechanisms for explicit timing in the sub-second range. *Neuroreport, 20*(10), 897-901. doi:10.1097/WNR.0b013e3283270b6e

Skagerlund, K., Karlsson, T., & Traff, U. (2016). Magnitude Processing in the Brain: An fMRI Study of Time, Space, and Numerosity as a Shared Cortical System. *Front Hum Neurosci, 10*, 500. doi:10.3389/fnhum.2016.00500

Smith, A., Taylor, E., Lidzba, K., & Rubia, K. (2003). A right hemispheric frontocerebellar network for time discrimination of several hundreds of milliseconds. *Neuroimage, 20*(1), 344-350. doi:10.1016/s1053-8119(03)00337-9

Smith, A. B., Giampietro, V., Brammer, M., Halari, R., Simmons, A., & Rubia, K. (2011). Functional development of fronto-striato-parietal networks associated with time perception. *Front Hum Neurosci, 5*, 136. doi:10.3389/fnhum.2011.00136

Taniwaki, T., Okayama, A., Yoshiura, T., Togao, O., Nakamura, Y., Yamasaki, T., . . . Tobimatsu, S. (2006). Functional network of the basal ganglia and cerebellar motor loops in vivo: different activation patterns between self-initiated and externally triggered movements. *Neuroimage, 31*(2), 745-753. doi:10.1016/j.neuroimage.2005.12.032

Teki, S., Grube, M., Kumar, S., & Griffiths, T. D. (2011). Distinct neural substrates of duration-based and beat-based auditory timing. *J Neurosci, 31*(10), 3805-3812. doi:10.1523/JNEUROSCI.5561-10.2011

Thaut, M. H., Demartin, M., & Sanes, J. N. (2008). Brain networks for integrative rhythm formation. *PLoS One, 3*(5), e2312. doi:10.1371/journal.pone.0002312

Tipples, J., Brattan, V., & Johnston, P. (2013). Neural bases for individual differences in the subjective experience of short durations (less than 2 seconds). *PLoS One, 8*(1), e54669. doi:10.1371/journal.pone.0054669

Tomasi, D., Wang, G. J., Studentsova, Y., & Volkow, N. D. (2015). Dissecting Neural Responses to Temporal Prediction, Attention, and Memory: Effects of Reward Learning and Interoception on Time Perception. *Cereb Cortex, 25*(10), 3856-3867. doi:10.1093/cercor/bhu269

Tracy, J. I., Faro, S. H., Mohamed, F. B., Pinsk, M., & Pinus, A. (2000). Functional localization of a "Time Keeper" function separate from attentional resources and task strategy. *Neuroimage, 11*(3), 228-242. doi:10.1006/nimg.2000.0535

Tregellas, J. R., Davalos, D. B., & Rojas, D. C. (2006). Effect of task difficulty on the functional anatomy of temporal processing. *Neuroimage, 32*(1), 307-315. doi:10.1016/j.neuroimage.2006.02.036

Ustun, S., Kale, E. H., & Cicek, M. (2017). Neural Networks for Time Perception and Working Memory. *Front Hum Neurosci, 11*, 83. doi:10.3389/fnhum.2017.00083

Wiener, M., Lee, Y. S., Lohoff, F. W., & Coslett, H. B. (2014). Individual differences in the morphometry and activation of time perception networks are influenced by dopamine genotype. *Neuroimage, 89*, 10-22. doi:10.1016/j.neuroimage.2013.11.019

Wittmann, M., Simmons, A. N., Aron, J. L., & Paulus, M. P. (2010). Accumulation of neural activity in the posterior insula encodes the passage of time. *Neuropsychologia, 48*(10), 3110-3120. doi:10.1016/j.neuropsychologia.2010.06.023

Wittmann, M., Simmons, A. N., Flagan, T., Lane, S. D., Wackermann, J., & Paulus, M. P. (2011). Neural substrates of time perception and impulsivity. *Brain Res, 1406*, 43-58. doi:10.1016/j.brainres.2011.06.048

Wittmann, M., van Wassenhove, V., Craig, A. D., & Paulus, M. P. (2010). The neural substrates of subjective time dilation. *Front Hum Neurosci, 4*, 2. doi:10.3389/neuro.09.002.2010

Woods, A. J., Hamilton, R. H., Kranjec, A., Minhaus, P., Bikson, M., Yu, J., & Chatterjee, A. (2014). Space, time, and causality in the human brain. *Neuroimage, 92*, 285-297. doi:10.1016/j.neuroimage.2014.02.015
