## Supplementary file E for "From *ATOM* to *GradiATOM*: Cortical gradients support time and space processing as revealed by a meta-analysis of neuroimaging studies"

**Supplementary Information E**

**Checklist for neuroimaging meta-analysis (Muller et al. 2017)**

| The research question is specifically defined | **YES** and it includes the following contrasts:  Aim 1:   - spatial processing > control conditions - temporal processing > control conditions   Aim 2:   - (spatial processing > control conditions) > (temporal processing > control conditions) - (spatial processing > control conditions) < (temporal processing > control conditions) |
| --- | --- |
| The literature search was systematic | **YES (PRISMA guidelines used)**, it includes the following keywords in the following databases:  **Keywords**:  For Space processing:  ((((((((mental rotation OR topographic OR (spatial AND (attention OR processing OR navigation OR memory) NOT time)) AND ((functional AND magnetic AND resonance AND imaging) OR fMRI)) NOT (structural OR DWI OR DTI OR diffusion OR machine learning OR multivariate OR multivoxel)) NOT (EEG OR transcranial OR stimulation OR tDCS))NOT (dementia OR deficit OR disorder OR pathology OR disease OR neglect OR cardia* OR Parkinson OR Alzheimer OR addiction OR drug OR psychiatric OR schizoph* OR psychosis OR neurologic OR injury OR stroke OR cannabis)) NOT (aging OR elderly OR children OR childhood)) NOT (mouse OR mice OR rat OR animal OR monkey)) NOT (meta-analysis[title] OR review[title] OR review[Publication Type] OR meta-analysis[Publication Type] OR Case Reports[Publication Type]))  For Time processing:  (((((((((time OR temporal) AND (duration OR processing OR perception) NOT (space OR spatial))) AND ((functional AND magnetic AND resonance AND imaging) OR fMRI)) NOT (structural OR DWI OR DTI OR diffusion OR machine learning OR multivariate OR multivoxel OR PET OR positron OR connectivity OR convolutional OR “deep learning”)) NOT (EEG OR transcranial OR stimulation OR tDCS)) NOT (dementia OR deficit OR disorder OR pathology OR disease OR neglect OR cardia* OR Parkinson OR Alzheimer OR addiction OR drug OR psychiatric OR schizoph* OR psychosis OR neurologic OR injury OR stroke OR cannabis)) NOT (aging OR elderly OR children OR childhood)) NOT (mouse OR mice OR rat OR animal OR monkey)) NOT (meta-analysis[title] OR review[title] OR review[Publication Type] OR meta-analysis[Publication Type] OR Case Reports[Publication Type]))  **Databases**: PubMed, MEDLINE, “related article” function in pubmed, reference within the selected literature. |
| Detailed inclusion and exclusion criteria are included | **YES**, and reason for non-standard criterion was:  Standard criteria applied: only whole brain experiment included; only studies reporting results in a standardized coordinate space were included.  Non standard criteria applied:   - studies that used fMRI or PET 🡪 criterion decided in order to maximize the power of the meta-analysis and in order to exclude studies with structural MRI; - studies analyzing the data using univariate approach that revealed localized increased activation were included 🡪 criterion decided to exclude papers that analyzed data using machine learning, whose results have a slightly different meaning; and to exclude papers using functional connectivity techniques, as we are not interested in connectivity; - studies with sample size of at least 5 participants (per group) were included 🡪 criterion decided according with previous coordinate based meta-analysis in order to reduce the likelihood to include studies presenting false positives; - studies were included only if they are performed on healthy individuals 🡪 criterion decided in accordance with the research question (to investigate the neural basis of normal spatial or temporal processing); - only studies reporting the contrast space or time > control condition were included 🡪 criterion decided to increase the specificity of the meta-analysis. - studies that did not focus on isolating specific brain regions’ activations related to a given spatial function (e.g., parahippocampal gyrus and posterior cingulate cortex) 🡪 criterion decided to highlight the activations linked to space processing rather than the activations associated with a specific spatial function (e.g., navigation or long-term memory) |
| Sample overlap was taken into account | **YES**, using the following method:  For each paper, only the contrast that most strongly reflects the process that the meta-analysis aims to investigate has been selected. In few cases, more contrasts were selected for a single paper: in all these cases, the authors run the analysis on two independent samples, and this is clearly stated in the paper. |
| All experiments use the same search coverage (state how brain coverage is assessed and how small volume corrections and conjunctions are taken into account) | **YES**, the search coverage is the following:  Whole brain.  If a study reported whole brain + ROI analysis, the whole brain analysis only has been included in the meta-analysis; if a study reported the ROI analysis only, the study was excluded from the meta-analysis in accordance with the inclusion/exclusion criteria.  We found no paper that applied small volume correction or hidden ROI. Papers reporting only conjunction analysis have been excluded. |
| Studies are converted to a common reference space | **YES**, using the following conversion:  Tailarach coordinates were reported into MNI space using a linear transformation as implemented in GingerALE 2.3.6 software. |
| Data extraction have been conducted by two investigators (ideal case) or double checked by the same investigator (state how double checking was performed) | **YES**, the following authors:  CS, NT (in the acknowledgements) checked inclusion criteria;  CS, NT (in the acknowledgements) extracted coordinates;  CS, NT (in the acknowledgements) extracted other info: Number of subjects included, Type of task (cognitive function involved); task (specific task); contrast performed; coordinate system; coordinate localization; associated statistic (t value, z score); p value criteria (corrected, uncorrected);  GC and MW double-checked the following data: extracted coordinates randomly and when discordant coordinates were extracted by CS and NT. |
| The paper includes a table with at least the references, basic study description (e.g. for fMRI task: stimuli), contrasts and basic sample descriptions (e.g. size, mean age and gender distribution, etc) of the included studies, source of information, reference space | **YES** and also the following data:  Study reference; Number of subjects included; task (specific task); contrast performed; coordinate system; coordinate localization; associated statistic (t value, z score); p value criteria (corrected, uncorrected);  **Please note that in this case the table is not enclosed within the paper but it is available within the supplementary information as an excel database.** |
| The study protocol was previously registered and all analyses planned beforehand, including the methods and parameters used for inference, correction for multiple testing, etc. | **NO**:   1) the meta-analysis was not registered before starting the search. Indeed, according with PROSPERO, systematic review and meta-analysis should be registered only if they are relevant to health and social care, which is not the case (https://www.crd.york.ac.uk/prospero/); 2) We declared that we planned all the analysis before starting the literature search and that we did not run any non-planned or non-prespecified analysis; 3) The meta-analysis used the default methods and parameters of the software with the following exceptions: none. |
| The meta-analysis includes diagnostics | NO. |
